# Somatic evolution of prostate cancer: mutation, selection, and epistasis across disease stages

**DOI:** 10.1101/2025.09.25.678387

**Authors:** Moein Rajaei, Alexander Yang, Kira A. Glasmacher, Christopher N. Cross, Nic Fisk, Elizabeth B. Perry, Jeffrey D. Mandell, Stephen G. Gaffney, Takafumi N. Yamaguchi, Julie Livingstone, Jose Costa, Peter Humphrey, Vincent L. Cannataro, Paul C. Boutros, Jeffrey P. Townsend

## Abstract

**Background:** Somatic mutations involved in prostate cancer tumorigenesis and disease progression have been identified, but their evolutionary dynamics—including differential selective pressures across oncogenesis and metastatic spread—remain poorly understood. No prior study has systematically quantified the adaptive landscape from prostate organogenesis through tumor initiation and progression to metastatic castrate-resistant prostate cancer (mCRPC), nor characterized the selective epistatic interactions that structure this evolutionary trajectory.

**Methods and Findings:** To address this gap, we analyzed 2,704 low- and high-risk primary tumors and metastatic castration-resistant prostate cancers to quantify the mutation rates, mutational processes, and scaled selection coefficients of somatic mutations across disease stages. Trinucleotide mutational patterns were stable, but both mutation load and mutation rates increased with progression. In parallel, selective pressures on specific somatic mutations changed substantially, revealing a dynamic adaptive landscape. Stage-specific selective effects were associated with significant synergistic and antagonistic selective epistasis among key driver genes. Early selection on *SPOP* mutations in the BRD3 binding domain were under strong positive selection, and they increased selection for subsequent *RHOA* mutations while decreasing selection for *TP53* mutations. Antagonistic selective epistasis was evident between mutations of *CUL3* and both *SPOP* and *PIK3CA*. Mutations in *KMT2C* increased the selection for mutations in *TP53*, consistent with their frequent co-occurrence. Synergistic epistatic interactions between mutations of *PTEN* and both *PIK3CA* and *AR* support a strong therapeutic rationale for combined inhibition of PI3K/AR pathway in PTEN-deficient prostate cancers.

**Conclusions:** These findings provide a comprehensive map of the evolving selective and epistatic forces that shape prostate cancer progression across clinical stages. By distinguishing shifts in selection from changes in mutation rate and revealing the extents of cooperative and conflicting relationships among driver mutations, our work identifies critical points of vulnerability and informs that design of therapeutic strategies that anticipate and intercept the somatic evolutionary trajectory of prostate cancer.

## Introduction

Prostate cancer is the second most diagnosed cancer in men worldwide, with over 1.4 million new cases and over 396,000 deaths reported in 2022 [1]. Incidence is predicted to double by 2040, reaching nearly three million new cases annually [2]. Early detection and treatment of localized disease substantially lowers the risk of metastasis and improves survival [3,4], whereas those who are diagnosed with distant metastases have a five-year survival of only 30% [5]. In many cancers, tumorigenesis has been described as a linear evolutionary progression from low-grade localized lesions to high-grade tumors and eventually to metastasis [6]. However, prostate cancer may diverge from this paradigm. Genomic studies suggest that low-grade and high-grade tumors have common precursors rather than undergoing direct evolutionary progression [7–10], and that low-grade and high-grade tumors diverge early [8,9,11–14]. Key driver mutations become prevalent among tumors at distinct stages, including at later stages during progression to advanced disease [15]. A comprehensive understanding of prostate cancer therefore requires analysis of how mutational patterns, gene-specific mutation rates, selective pressures, and epistatic interactions among driver genes vary across grades and stages of disease.

Comprehensive genomic profiling has revealed the complex genetic heterogeneity of prostate cancer, marked by recurrent alterations in multiple key pathways. Frequently mutated driver genes include *SPOP, TP53, MED12* and *FOXA1*; as well as alterations of the androgen receptor (*AR*). Defects in DNA damage repair pathways involving *BRCA* and *ATM* and gene fusions involving ETS family genes are fairly common [12,16–28]. The Wnt signaling pathway is often altered in metastatic castration-resistant prostate cancer (mCRPC), with mutations in *CTNNB1, SMAD4*, and *APC* [17,29,30], and is also implicated in aggressive localized disease through alterations in genes such as *BRCA2* and *ZNRF3* [12,31]. The PI3K pathway is commonly affected *via* mutations and copy number alterations (CNAs) in *PIK3CA* and *PIK3CB* as well as *AKT1* and *PTEN [32–34]*. mCRPC is further characterized by both gain-of-function alterations of *AR* or regulators of *AR* and its upstream regulator *FOXA1* [16], as well as loss-of-function mutations in AR pathway repressors [35,36]. Aberrant post-translational modifications and alternative splicing of AR can also promote ligand-independent activity, driving tumor progression despite androgen deprivation [37,38]. However, the evolutionary dynamics that drive progression from prostate organogenesis to low-grade and high-grade localized cancer—and ultimately to mCRPC—remain poorly understood. Previous studies have identified statistically significant differences in prevalence of somatic variants between primary and mCRPC tumors [31,39] and have constructed prostate cancer progression and tumor evolution models based on conditional probabilities of variant co-occurrence [40,41]. These approaches rely on prevalences and *P* values, which are not direct measures of cancer effect. Prevalence reflects the proportion of tumors harboring a specific variant, and *P* values quantify statistical confidence rather than biological effect [42]. Quantifying the strength of selection—a true effect size—provides a biologically grounded basis for prioritizing driver mutations in basic research, identifying therapeutic targets, guiding drug development, and designing biomarker-driven clinical trials [43].

Accordingly, prevalence-based measures of co-occurrence and mutual exclusivity among mutations can be confounded by shared or divergent mutagenic processes across tumors, masking actual functional interactions or falsely indicating their presence [44]. In contrast, inference of selective epistasis—defined as deviations in the combined selective effect of mutations from the product of their independent selective effects—offers a direct measure of fitness interactions between driver genes [45]. No prior study has systematically quantified the adaptive landscape from organogenesis through tumorigenesis and metastatic progression, nor characterized the selective epistatic interactions that structure this evolutionary trajectory.

Here we characterize the evolutionary dynamics of somatic mutation and selection that drive prostate cancer development and progression from low-grade and high-grade localized tumors to mCRPC. Applying a somatic evolutionary framework, we quantify mutation rates and cancer effect sizes, which estimate the selective advantage conferred by each mutation in the context of tumor evolution [42]. By analyzing mutations in their pairwise context, incorporating underlying mutation rates and correlations, we identify the time-ordered selective interdependence of somatic mutations that occupy key roles in the development of prostate cancer [42]. These analyses reveal the shifting selective landscape across tumor grades and stages, providing a map of the mutational trajectories and epistatic interactions that underline prostate cancer evolution.

## Methods

### Study design, dataset assembly and analysis

We assembled a dataset of 2,704 tumors, including 1,122 whole-exome sequences (WES) and 1,289 targeted gene sequences (TGS) from three sources [26,46,47], as well as 273 whole-genome sequences (WGS) of primary prostate tumors provided by Espiritu *et al*. [15], and 20 WGS from TCGA. Samples were excluded if they contained only nucleotide base substitutions at known germline variant sites, within repetitive regions, or no nucleotide base substitutions, yielding a final set of 2,618 samples for downstream analysis, comprising 1,593 primary and 1,025 mCRPC tumors. Primary tumors were stratified by pathologic grade: Grade Groups 1 or 2 were classified as low-grade (*n* = 479), and Grade Groups of 3, 4, or 5 were classified as high-grade (*n* = 406) [48], based on findings that Grade-Group 1/2 tumors and Grade Group 3/4/5 have very different risks of biochemical recurrence after definitive local therapy [49]. Grade Group data were unavailable for 708 primary tumors, which were omitted from grade-dependent analyses. Targeted sequence data was gathered using panels covering 341, 410, and 468 cancer-relevant genes.

### Step-specific mutation rates from organogenesis to grade-group primary tumors and from grade-group primary tumors to mCRPC

To investigate the step-specific rates at which cancer cells are mutated in specific genes, we partitioned the tumor samples by pathologic grade and disease progression. We modeled two key steps: step 0→1 refers to the evolutionary step from prostate organogenesis to primary tumor formation (either low-grade or high-grade), for which we have directly applicable data from primary tumor samples, and step 1→2, on which we don’t have directly applicable data—instead, we have data on the results of the full trajectory 0→2 denoting the sum of mutations accumulated through the first step (from prostate organogenesis to primary tumors) and the second step (from either low-grade or high-grade primary tumors to mCRPC). To quantify gene mutation rates (i) from prostate organogenesis to low-grade primary tumors (*U* _*L*,0→1,*g*_), (ii) from prostate organogenesis to high-grade primary tumors (*U* _*H*,0→1,*g*_), and (iii) from prostate organogenesis through primary progression to mCRPC (*U*_0→2,*g*_), we used the dNdScv; RRID:SCR_017093 model [50] for synonymous-site, covariate-adjusted mutation rate estimation, implemented via cancereffectsizeR v2.7.0; [51 v. 2.7.0] [51]. We supplied the prostate tissue-specific covariates previously described in Cannataro et al. [42]. We divided the maximum-likelihood expected number of synonymous substitutions in the cohort by the number of samples in respective cohorts, then by the number of synonymous sites in each gene to yield the three expected rates of mutation per nucleotide for each gene *g*: *U* _*L*,0→1,*g*_, *U* _*H*,0→1,*g*_, and *U*_0→2,*g*_ . The proportion of mutations accumulated in the steps from prostate organogenesis to low-grade and prostate organogenesis to high-grade primary tumors for each gene was assessed separately as *W*_0→1,*g*_ = *U*_0→1,*g*_ /*U*_0→2,*g*_, and the proportion accumulated in the steps from low-grade primary tumors to mCRPC and high-grade primary tumors to mCRPC was *W*_1→2,*g*_ = 1 − *W*_0→1,*g*_ .

For tumors with ≥ 50 SNVs, we estimated trinucleotide-context-specific mutation rates using COSMIC v3.2 mutational signature extraction via MutationalPatterns v3.8.1; RRID:SCR_024247 [52], fitting a set of signatures associated with mutational processes active in prostate cancer as in Cannataro *et al* [53]. Since targeted gene sequencing data contain too few mutations and are biased toward cancer hotspots they were excluded from the mutation rate calculation; average mutation rates for each site across whole-exome sequenced samples were assigned for all targeted gene sequence data.

### Mutation rates for epistatic selection

To evaluate pairwise selective epistasis between somatic mutations along the full trajectory from organogenesis to mCRPC, we analyzed all samples regardless of stage. Sample-specific site mutation rates were calculated with the standard cancereffectsizeR workflow [51], without partitioning mutation rates as in the stepwise selection model.

### Scaled selection coefficients from organogenesis to grade-group primary tumors and from grade-group primary tumors to mCRPC: estimation of step-specific cancer effect sizes

We used cancereffectsizeR [51 v. 2.7.0] to calculate sample-specific mutation rates. For each tumor *i*, we calculated a mutation rate *µ*_0→2,*g,i*_ by site-specific sample-specific patterns of mutation obtained *via* mutational signature analysis [42], summed over all variants in a gene *g*. Then we defined the mutation rate for the defined set of sites during the initial step as *µ*_0→1,*g,i*_ = *W*_0→1,*g*_ × *µ*_0→2,*g,i*_, and during the corresponding second step as *µ*_1→2,*g,i*_ = *W*_1→2,*g*_ × *µ*_0→2,*g,i*_ .

We quantified step-specific strengths of selection on SNVs and genes with cancereffectsizeR v. 2.7.0 [51 v. 2.7.0], using hg19 reference coordinates. We focused our analysis on 16 genes that are well established as drivers of tumorigenesis, most of them established as specific drivers of prostate cancer [16,17,20,22,32,33,54–56]. For our step-specific analyses and estimates of epistatic effects, strengths of selection were quantified for sets of driver sites across the gene rather than for each amino acid. We maximized our power to observe differences in selection pressure by maximizing the likelihood of a single strength of selection that operated on all intragenic variant sites within our dataset.

In Cannataro *et al*. [53,57], likelihood for the strength of selection *γ* was calculated for any single cohort of tumors by maximizing

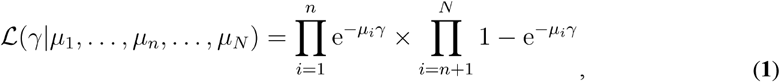

where *µ*_*i*_,1 ≤*i* ≤*N*, is the mutation rate of a defined set of sites in sample *i*, and where *n* < *N* are defined such that any variant in the defined set of sites is absent in *n* samples and is present in samples *n*+1 …*N*, and where the subscript *g* is suppressed in **Eq. 1** and later equations for clarity of presentation.

To estimate scaled selection coefficients along a multi-step sequential model of tumorigenesis and metastasis, we extended **Eq. 1** to differentiate two steps—evolution from prostate organogenesis to primary tumors (Step 0→1) and evolution from primary tumors to mCRPC (Step 1→2):

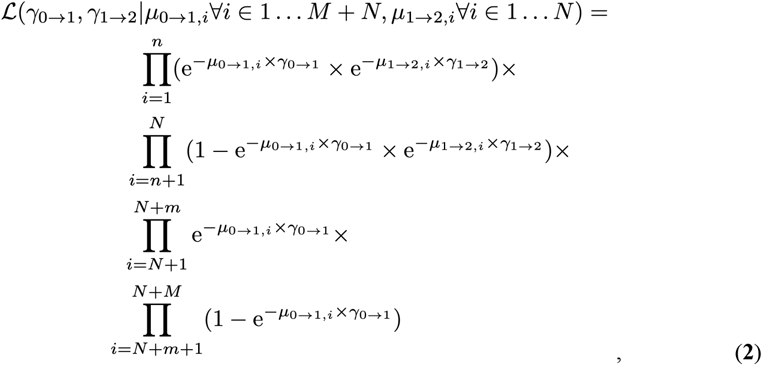

for a dataset with *M* primary tumor tissue samples, of which *M* − *m* exhibit a substitution and *m* do not, and where there are *N* mCRPC tissue samples in total, of which *N* − *n* exhibit a substitution and *n* do not. The primary tumor samples are identified by index *i* =1+*N*, …,*m* +*N*, … *M* +*N* and the mCRPC samples are also identified by index *i* = 1,…,*n*, …,*N*. The first product term in **Eq. 2** represents mCRPC samples where all variants in the defined set of sites are absent, the second product term represents mCRPC samples where any such variant is present, the third product represents primary tumor samples where all such variants are absent, and the fourth product represents primary tumor samples with any such variant present.

**Eq. 2** can further be generalized for any number of steps *S*:

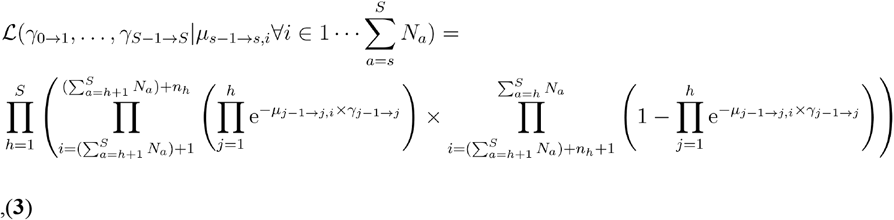

where *s* = 1, …,*S* denotes the step of progression toward cancer, and *S* corresponds to the last step present in the data. For each step *s*, there are *N*_*s*_ samples: *n*_*s*_ without a variant and *N*_*s*_−*n*_*s*_ with a variant in the defined set of sites. The product index *h* specifies the step of a sample, *i* specifies the sample index, and *j* functions as an index that breaks down all *h* steps of a sample’s trajectory into the distinct stepwise processes. For each step, the samples are indexed by

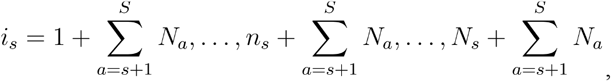

resulting in

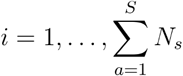

across all steps.

As part of a separate analysis of robustness of results to model of progression, we also performed this gene-specific mutation rate estimation on a linear three-step evolution model (**Eq. 3**) implemented in cancereffectsizeR, where mutations occurred (i) from prostate organogenesis to low-grade primary tumors, (ii) from low-grade to high-grade primary tumors, and (iii) from high-grade primary tumors to mCRPC.

### Protein structural modeling and visualization

The structural model for the Speckle-type POZ protein SPOP was sourced from the Research Collaboratory for Structural Bioinformatics Protein Data Bank (UID: 6I41; RRID:SCR_012820) [58]. The structural model for the ligand-binding domain of the androgen receptor AR was sourced from the RCSB PDB as well (UID:3L3X) [59]. The protein structure was then visualized, and important amino-acid residues were highlighted using the UCSF Chimera software; RRID:SCR_004097 [60].

### Statistical analysis of selective epistasis among driver genes

The positive or negative change in selection pressure imposed on a new variant in one gene by the presence of an oncogenic variant in another gene was estimated using the analytical solution for pairwise epistasis inference derived in Alfaro-Murillo & Townsend [45] implemented in cancereffectsizeR [42]. For any pair of variants, the likelihood of sample genotypes was calculated as a product of probabilities of serial fixation based on neutral mutation rates and somatic genotype-specific strengths of selection. Likelihoods were maximized to estimate unknown selection parameters. Univariate confidence intervals for each parameter were estimated by fixing the other parameters at their maximum likelihood estimates, assuming the likelihood is close to its asymptotic chi-square form and calculating the univariate parameter values that reduce the optimal likelihood by χ2(0.05, 1) / 2 = 1.92. To maximize statistical power to discern epistatic effects, we conducted all analyses of epistatic effects enforcing inference of a single strength of selection for all recurrent variants within each gene, except for *ROCK1*, which was excluded due to its heterogeneity of coverage in sequencing panels.

## Results

### Prevalence of driver mutations varies among low-grade, high-grade, and mCRPC

We began by characterizing the prevalence of somatic mutations in key driver genes across low-grade, high-grade, and metastatic castration-resistant prostate cancer (mCRPC) tumors (**Fig 1**). In low-grade tumors, mutations in *SPOP* were more prevalent than in any other drivers. In high-grade tumors, somatic mutations in *SPOP* and *TP53* genes were more prevalent than in any other drivers. In mCRPC, *TP53* and *AR* mutations exhibited the highest prevalence; *AR* mutations were substantially more prevalent in mCRPC than in both low-grade and high-grade tumors (**Fig 1**). High prevalence in low-grade, high-grade, or mCRPC indicates a large patient population, an important metric of how many people might benefit if these mutations prove to be both oncogenically important and targetable. However, prevalence alone does not measure the cancer effect or oncogenic impact of the mutation. To quantify cancer effects, underlying mutation rates that substantially impact occurrence of variants must be deconvolved from the prevalence.

**Fig 1.**
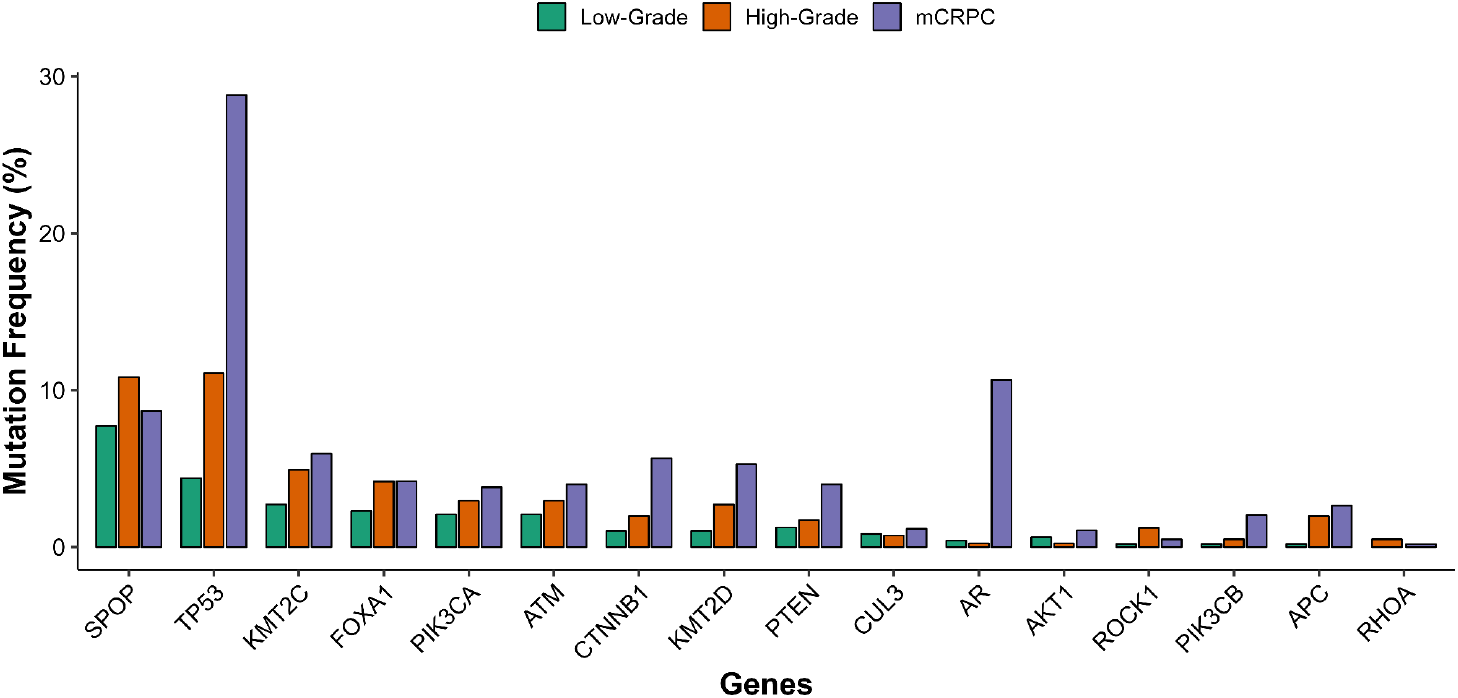
Prevalence of variants in 16 selected driver genes, in low-grade primary tumors (*n* = 479), high-grade primary tumors *(n* = 406), and metastatic castration-resistant prostate cancer (*n* = 1,025).

### Trinucleotide mutation signatures and gene mutation rates are similar in low-grade, high-grade, and mCRPC

To determine whether differences in driver mutation prevalence across prostate cancer stages could be attributed to distinct mutational processes, we analyzed trinucleotide mutation signatures from prostate organogenesis to low-grade tumors, from organogenesis to high-grade tumors, and from primary tumors to mCRPC. Across these somatic evolutionary trajectories, the cosine similarity of trinucleotide mutation spectra was high (low-grade vs high-grade: 0.97, low-grade vs mCRPC: 0.80, high-grade vs mCRPC: 0.91; **Fig 2**). These similarities persisted in analyses restricted to an age-matched cohort (**Fig S1**), indicating that age-related differences did not confound the result.

**Fig 2.**
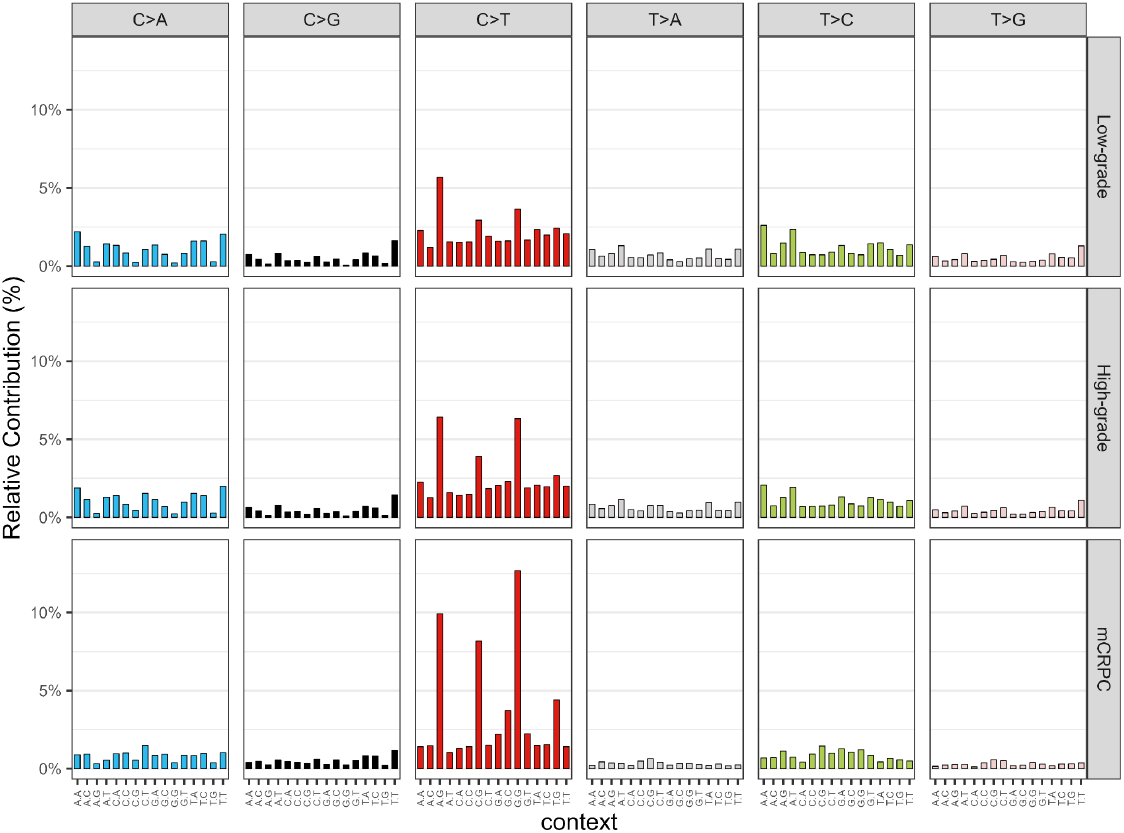
Percent of single-nucleotide somatic variants within each trinucleotide context in low-grade primary tumors, high-grade primary tumors, and metastatic castrate-resistant prostate cancer.

Correspondingly, all three tumor groups exhibit the same rank order of contribution of the major COSMIC v3.2 mutational signature weights [61] contributing to mutagenesis in prostate cancer (**Fig S2**). Age-associated clock-like signatures SBS1, SBS5, and SBS40 predominated in samples, as has been previously reported in prostate cancer [62]. A marked enrichment in C→T transitions throughout the whole-genome and whole-exome sequenced tumors, especially in the ACG/CCG/GCG trinucleotide contexts, reflected SBS1 activity, associated with spontaneous or enzymatic deamination of 5-methylcytosine to thymine. Enrichment of T→C mutations in the ATA/ATG/ATT contexts aligns with SBS5. SBS1 and SBS5 are clock-like signatures that correlate with the patient’s age [53]—and age is a significant contributor to oncogenic mutations in prostate cancer [53] and a significant risk factor to developing prostate cancer [63].

Unlike other cancers with strong exogenous mutagenic exposures, such as melanoma, lung cancer, or hepatocellular carcinoma, prostate cancer tissues are riddled with mutational patterns attributable to age-related mutational processes. These largely stable mutational signatures suggest that differences in driver gene prevalence are not driven by changes in mechanism of mutation.

Tumor mutational burden is elevated in mCRPC compared to primary tumors, as reported previously [16,17,24,64]. However, tumor mutational burden is a product over time of underlying mutation rates and selection for the mutations induced. To dissect these contributions, we compared gene-specific neutral mutation rates along the steps from prostate organogenesis to low-grade and high-grade primary tumors and the steps from prostate organogenesis to mCRPC (**Fig 3**).

**Fig 3.**
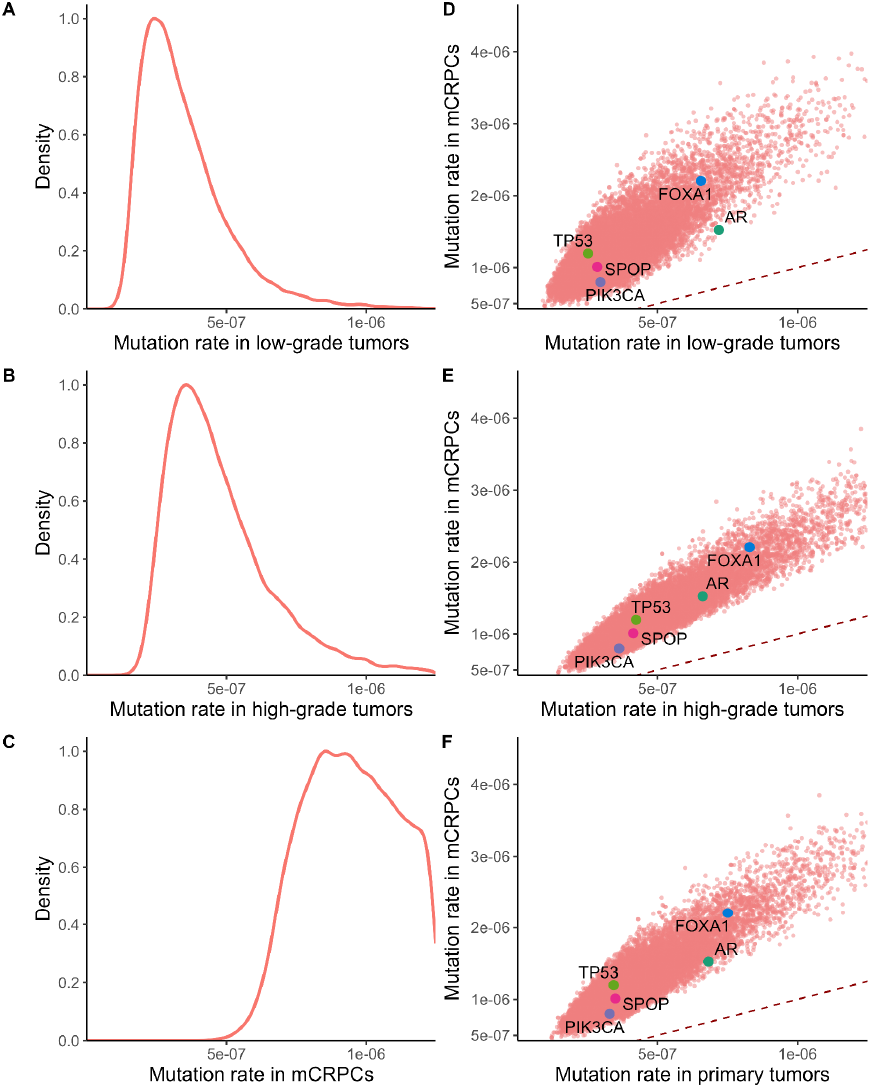
The gene-level mutation rates spanning from (**A**) organogenesis to low-grade primary tumors, (**B**) organogenesis to high-grade primary tumors, (**C**) organogenesis to metastatic castrate-resistant prostate cancer, and in (**D**) organogenesis to metastatic castrate-resistant prostate cancer versus organogenesis to low-grade primary tumorigenesis, (**E**) organogenesis to metastatic castrate-resistant prostate cancer versus organogenesis to high-grade primary tumorigenesis, (**F**) organogenesis to metastatic castrate-resistant prostate cancer versus primary tumorigenesis. For reference, equality is graphed (dashed red line), and five well-characterized prostate cancer drivers *SPOP* (pink), *PI3KCA* (dark blue), *TP53* (light green), *FOXA1* (blue), and *AR* (green) are indicated.

The rate at which neutral mutations are introduced into each gene were significantly higher along the full trajectory from organogenesis to mCRPC than along the steps from prostate organogenesis to low-grade tumors (rank-biserial correlation: -0.85; *P* < 2.2 × 10^−16^; Mann-Whitney U test; **Fig 3D**) or to high-grade tumors (rank-biserial correlation: -0.81, *P* < 2.2 × 10^−16^, Mann-Whitney U test; **Fig 3E**). When comparing the organogenesis-to-mCRPC trajectory with the entire graded primary tumor cohort (low- and high-grade combined), neutral rates again increased significantly (including both low- and high-grade; rank-biserial correlation: 0.83, *P* < 2.2 × 10^−16^, Mann-Whitney U test; **Fig 3F**).

Mutation rates varied substantially across genes—even among well-characterized prostate cancer driver genes such as *SPOP, TP53, PIK3CA, FOXA1*, and *AR* (**Fig 3D–F and S1 Table**) that play distinct roles in development, progression, or treatment resistance [22,25]. All 16 genes we identified as well-established drivers of prostate cancer [16,17,20,22,32,54–56] exhibited a higher neutral mutation rate along the trajectory from organogenesis to mCRPC than during earlier steps toward primary tumors (**Fig 3D–F and S1 Table**). For example, neutral mutation rates of the androgen-receptor gene *AR*, the oncogene *FOXA1*, and tumor-suppressor gene *TP53* were markedly elevated during progression from primary tumors to mCRPC. These findings connect increasing endogenous mutational input—linked to changes in cell state or tumor context—to elevated tumor mutational burden and evolutionary plasticity observed in mCRPC. Because driver genes exhibit distinct (and typically increasing) mutation rates over the course of disease, their prevalence at different developmental steps cannot be directly interpreted as evidence of selective advantage. Accurate inference of selective advantage requires use of a model that accounts for these gene- and step-specific differences in mutation rate.

### Mutations of prostate cancer driver genes are subject to markedly different selection in low-grade, high-grade, and mCRPC tumors

Stepwise analysis of selection intensity revealed that the differences in driver mutation prevalence across distinct steps of prostate cancer cannot be attributed solely to changing gene-specific mutation rates. Instead, the selection operating on novel mutations varies substantially over the course of disease progression (**Fig 4**). Among the 16 key driver genes analyzed, *SPOP, AKT1, KMT2D, CTNNB1, CUL3, PIK3CA, TP53, FOXA1, ATM*, and *KMT2C* were under strong selection during the step from prostate organogenesis to low-grade tumors, with markedly reduced selection during progression to mCRPC. *PTEN* mutations were strongly selected during tumor initiation in both low-grade and high-grade primary tumors but showed diminished selection intensity during metastatic progression. In contrast, mutations in *APC, ROCK1, RHOA*, and *PIK3CB* were more strongly selected during the steps from prostate organogenesis to high-grade prostate tumors, associating these genes with aggressive primary disease. Mutations in *AR* were strongly selected during the steps from primary tumors to mCRPC, reflecting the role of the androgen receptor in the evolution of resistance to antiandrogenic therapies prior to mCRPC [16,65].

**Fig 4.**
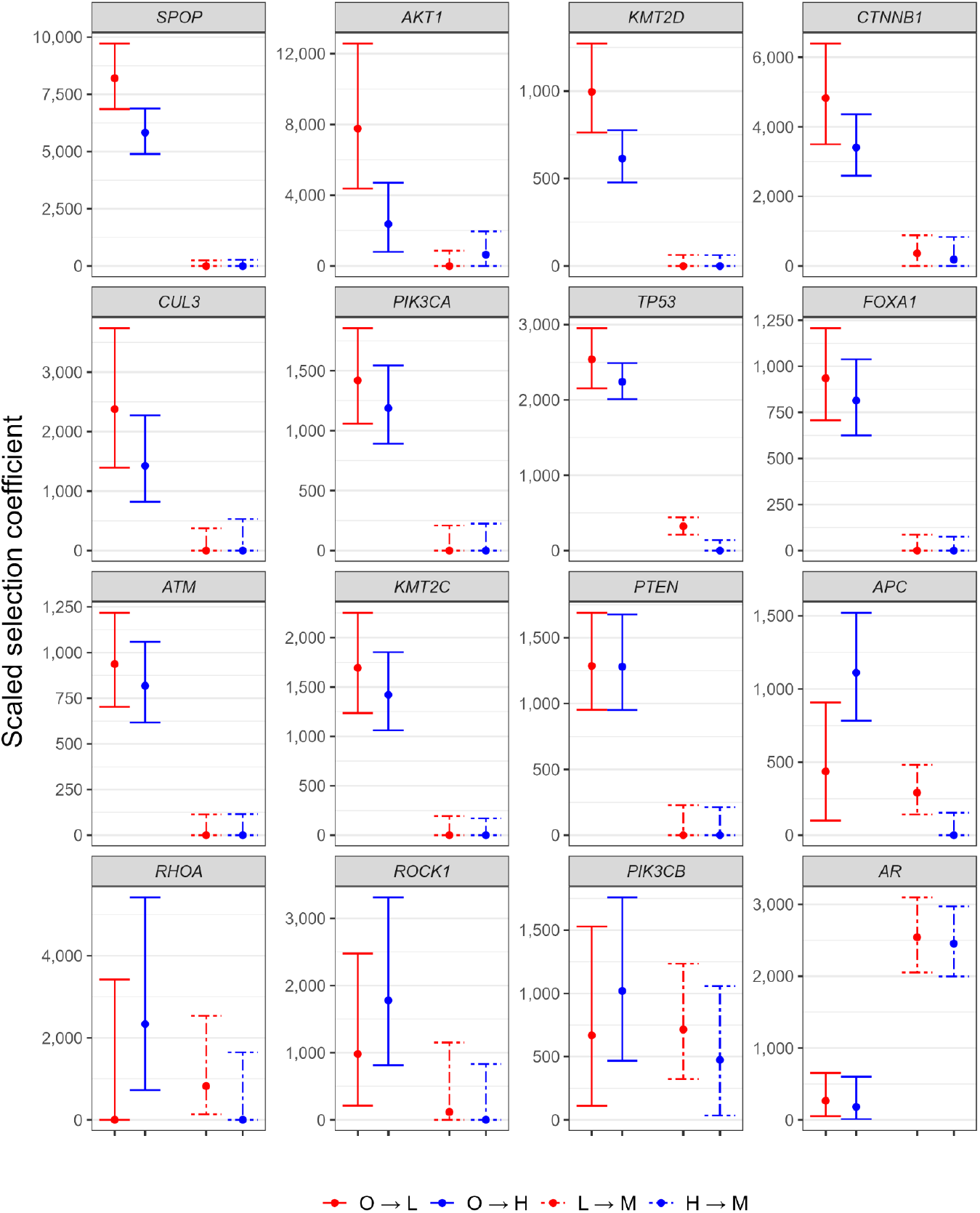
Gene-level estimates and 95% confidence intervals for scaled selection coefficients on somatic variants in oncogenic sites of 16 genes that are known to act as drivers in prostate cancer tumorigenesis and metastasis, from prostate organogenesis to low-grade primary tumors (O→L, red), from prostate organogenesis to high-grade primary tumors (O→H, blue), from low-grade tumors to metastatic castrate-resistant prostate cancer (L→M, red dot-dashed), and from high-grade tumors to metastatic castrate-resistant prostate cancer (H→M, blue dot-dashed).

These patterns of selection were robust to how prostate cancer was modeled—whether as distinct somatic genetic trajectories through low-grade and high-grade prostate cancer to mCRPC [11], or as a single trajectory from low-grade to high-grade prostate cancer to mCRPC. Under this three-step serial model, strong positive selection was again inferred for *SPOP, AKT1, KMT2D, CTNNB1, CUL3, PIK3CA, TP53, FOXA1, ATM*, and *KMT2C* during progression from prostate organogenesis to low-grade prostate cancer (**Fig S3**). During subsequent progression from low-grade to high-grade disease, mutations in *APC, ROCK1, RHOA*, and *PIK3CB* experienced strong selection, consistent with their involvement in the emergence of more aggressive localized disease [55,66]. Finally, mutation rates inferred for *AR* were not consistent with a three-step model; however, based on a two-step organogenesis-to-primary-to-mCRPC model, AR mutations yielded strong selection on new mutations exclusively during the step from primary tumors to mCRPC, underscoring their central role in late-stage, treatment-resistant disease.

### Variant-specific selection in SPOP was associated with structural disruption of BRD3 binding

For genes with multiple recurrent mutations, such as *SPOP* and *AR*, scaled selection coefficients can be estimated for individual amino-acid variants. Across the full cohort of prostate cancer samples, numerous *SPOP* amino-acid sites were mutated (**Fig 5**). Among these sites, nine were recurrently mutated: three sites, S119N, K129E, and K134N, that exhibited only one recurrent substitution, and six sites, Y87•, F102•, F125•, D130•, F133•, and W131• that exhibited multiple distinct substitutions.

**Fig 5.**
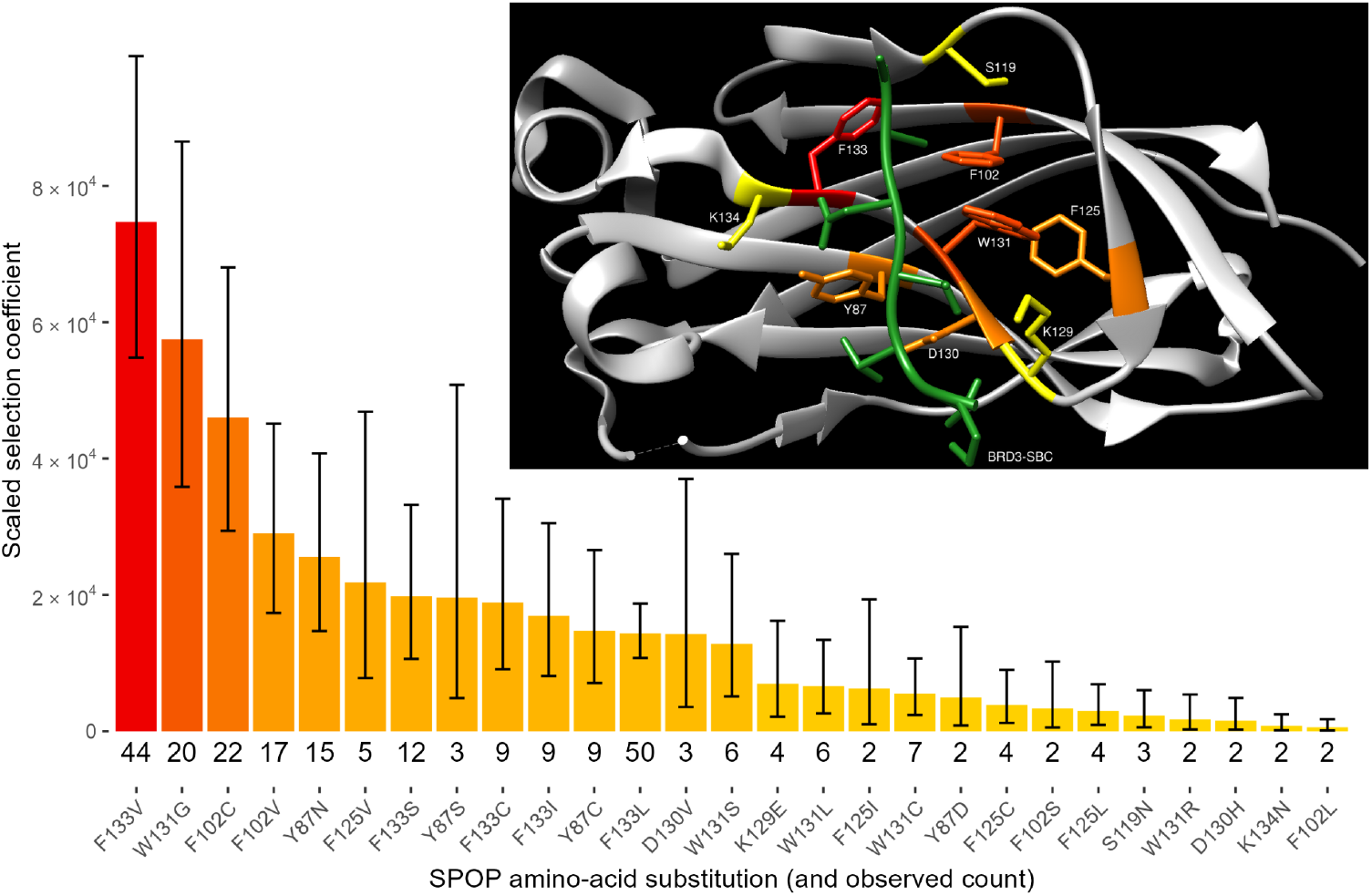
Scaled selection coefficients for recurrent single-nucleotide variant amino-acid substitutions in *SPOP* during the evolutionary trajectory from prostate organogenesis to primary and metastatic castrate-resistant prostate cancer tumors, ranging from strongly selected (red) to weaker-selected (yellow), as well as 95% confidence intervals for the scaled strength of selection and observed counts of variants out of *n* = 2,618 tumors with sequence data. *Inset*: *SPOP* protein structure (grey), with the binding moiety of the *SPOP* ligand, BRD3 (green); for illustrative purposes, all substituted sites (red: stronger selection to yellow: weaker selection) are depicted simultaneously in a composite structure, featuring for each site the strongest-selected recurrent amino-acid substitution.

Examination of the structural location of these mutations and the amino-acid substitutions they produce reveals why certain somatic variants exhibit higher scaled selection coefficients than others. Mutated residues either directly form the substrate-binding cleft (Y87•, F102•, K129•, D130•, F133•, W131•; average scaled selection coefficient 1. 9 × 10^4^) or are immediately adjacent to it (S119•, F125•, K134•; average scaled selection coefficient 6. 3 × 10^3^) [67]. Among all variants, SPOP F133V and SPOP W131G exhibit the highest scaled selection coefficients.

The selective advantage of SPOP F133V is statistically significantly greater than all but two other mutations (W131G and F102C; non-overlapping 95% CIs.) These amino-acid changes constitute striking physiochemical and structural alterations, reducing two bulky aromatic or hydrophobic residues (W→G and F→V) to much smaller R-groups, at structurally critical sites. These changes likely disrupt SPOP-substrate interactions and impair ubiquitin-mediated degradation, facilitating oncogenic signaling and driving clonal expansion. Notably, the most prevalent variant, SPOP F133L—detected in 50 tumors—ranks only 12^th^ by scaled selection coefficient. This discordance between prevalence and scaled selection coefficient is a consequence of an exceptionally high mutation rate at the site, attributable to a highly mutable trinucleotide context [68] and the fact that this specific missense mutation can result from three distinct SNVs. These findings underscore the variable impacts of distinct *SPOP* variants on prostate cancer oncogenesis.

### Variant-specific selection and functional impacts of AR variants in its ligand-binding domain

Overactivation of the *AR* signaling pathway is a hallmark of mCRPCs, driven by point mutations, gene amplifications, and alternative splice variants [69,70]. In our cohort, we identified nine distinct recurrent point mutations affecting seven codons in *AR* (**Fig 6**), all localized to its ligand-binding domain (LBD; **Fig 6**, *inset*) [71]. Indeed, the four most strongly selected mutations—L702H, T878A, H875Y, and W742C—were collectively present in 10% of all mCRPCs (**Fig 6**) [17]. Despite their shared localization within the LBD, these mutations exhibited a broad range of selective effects (**Fig 6**). L702H conferred the greatest selective advantage, followed by T878A and H875Y. Two distinct mutations at codon 742 (W→L and W→C) were indistinguishable in effect. Functionally, T878A, H875Y, W742C, and W742L point mutations are well-characterized activating mutations that enable aberrant activation of the AR protein in response to first-generation anti-androgens such as flutamide and bicalutamide, transforming these therapies from antagonists into partial agonists and promoting continued AR signaling despite therapeutic blockade [72–75].

**Fig 6.**
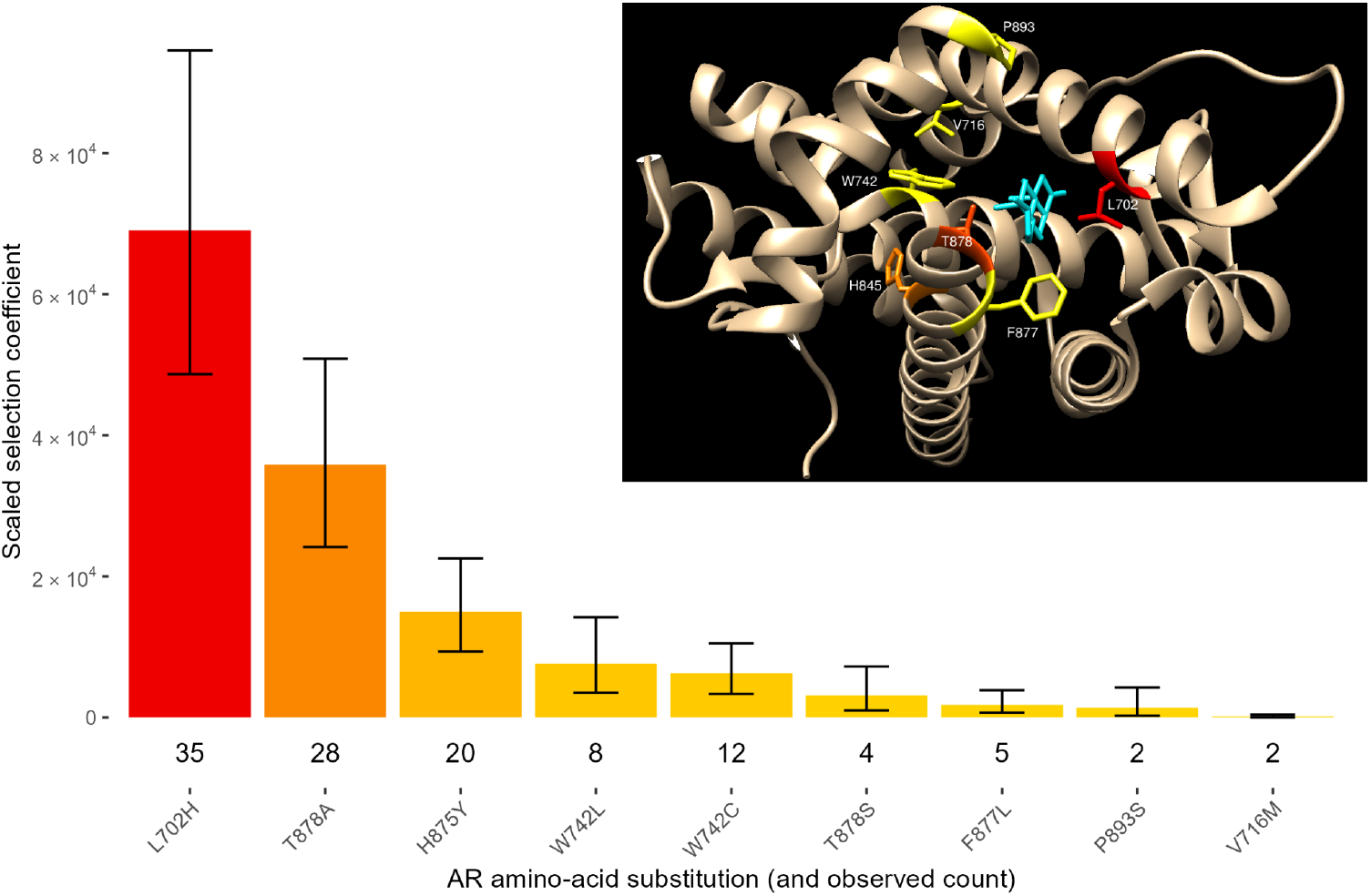
Scaled selection coefficients of recurrent single-nucleotide variant amino-acid substitutions in *AR* along the step from primary tumors to metastatic castrate-resistant prostate cancer, ranging from strongly selected (red) to weaker-selected (yellow), 95% confidence intervals for the scaled strength of selection, and observed counts of variants out of *n* = 1,025 tumors with sequence data. *Inset*: AR protein structure (grey), with the binding moiety of the AR ligand, dihydrotestosterone (teal); all substituted sites (stronger selection:red to weaker selection:yellow) are depicted simultaneously in a composite structure, featuring for each site the strongest-selected recurrent amino-acid substitution. The residues spanning S760 to M781 were removed from the image to provide a better perspective view into the ligand-binding pocket.

Unlike anti-androgen sensitizing mutations, L702H activates the androgen receptor through a distinct mechanism: it confers sensitivity to endogenous glucocorticoids, enabling androgen-independent transcriptional activity [76,77]. This heterogeneity of mechanism reinforces the central importance of the androgen receptor signaling axis in prostate cancer tumorigenesis. Moreover, the L702H is under substantially stronger positive selection—nearly twice that of T878A, the most strongly selected anti-androgen sensitizing mutation, and fourfold or more greater than others—quantifying its markedly greater proliferative or survival advantage. This strong selection for the L702H mutation indicates that glucocorticoid-driven AR activation represents a potent route to therapeutic resistance in mCRPC, and provides a compelling rationale for the development of mechanism-specific therapeutic strategies designed to counteract AR reactivation by glucocorticoids in patients with L702H-mutated tumors.

### Analysis of selective epistasis among driver genes indicates somatic evolutionary chronology of prostate cancer

Pairwise epistasis analysis of 15 prostate cancer driver genes across tumors provided insights into how oncogenic variants in one gene influence the selection for variants in the others. Of the 210 possible pairwise interactions among these genes, 36 exhibited significant synergistic epistasis. Many of these interactions aligned with the evolution of low-grade and high-grade prostate cancer, along with their progression to mCRPC (**S2 Table**).

This analysis also revealed asymmetric interactions and time-ordered dependencies that clarify the evolutionary trajectory of prostate cancer tumorigenesis. For instance, the trend for mutations in most driver genes (10 out of 14) was to reduce selection for mutations in *SPOP* (**Fig 7A**), with notable exceptions. Mutations in *RHOA, PIK3CA, AR*, and *CTNNB1* trended toward positive interactions with *SPOP* mutations, whereas *SPOP* mutations trended toward increasing the selection for *AKT1* and strongly increasing selection for *RHOA*. Substitutions in *SPOP* play an early, initiating role in prostate cancer tumorigenesis, enhancing the degree of proliferation and survival attributable to subsequent mutations that are promoting progression.

**Fig 7.**
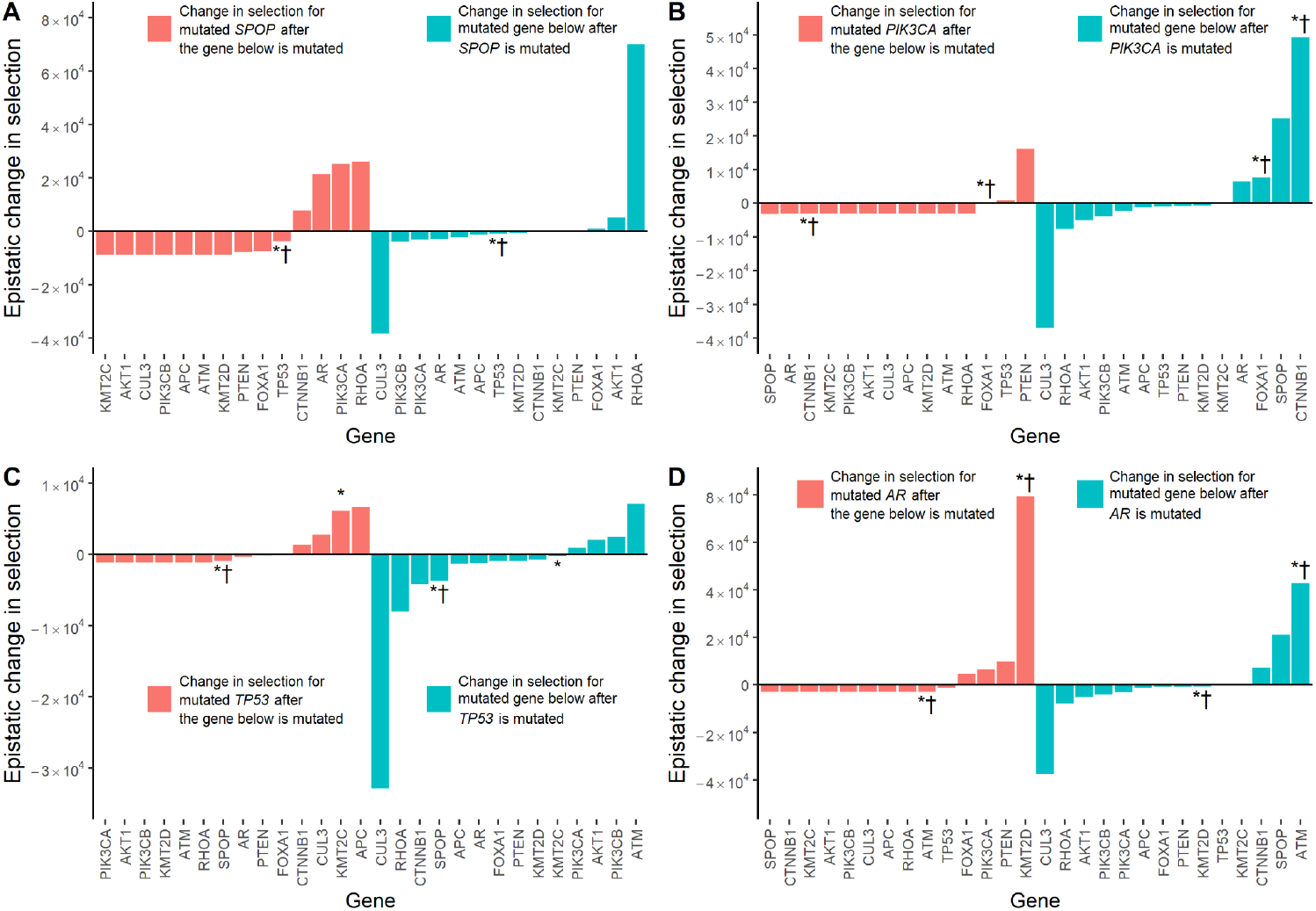
Pairwise epistatic effect trends of 15 genes based on all 2,699 tumors, focusing on *SPOP, PIK3CA, TP53*, and *AR*. (**A**) Change in scaled selection coefficient for mutated *SPOP* due to epistatic effects of mutations in 14 genes (orange), and change in scaled selection coefficient for mutations due to epistatic effects of mutated *SPOP* (aquamarine). (**B**) Change in scaled selection coefficient for mutated *PIK3CA* due to epistatic effects of mutations in 14 genes (orange), and change in scaled selection coefficient for mutations due to epistatic effects of mutated *PIK3CA* (aquamarine). (**C**) Change in scaled selection coefficient for mutated *TP53* due to epistatic effects of mutations in 14 genes (orange), and change in scaled selection coefficient for mutation due to epistatic effects of mutated *TP53* (aquamarine). (**D**) Change in scaled selection coefficient for mutated *AR* due to epistatic effects of mutations in 14 genes (orange), and change in scaled selection coefficient for mutations due to epistatic effects of mutated *AR* (aquamarine). Symbols at the end of each bar indicate statistical significance for mutual exclusivity or co-occurrence (*: Fisher’s exact test, *P* < 0.05) and for non-directional epistasis (†: cancereffectsizeR, *P* < 0.05).

*PIK3CA* mutations exhibit moderate positive interactions with earlier driver mutations, such as *SPOP*. Mutations in *PIK3CA* experience enhanced selection following substitutions in *PTEN* and to a lesser degree *TP53* (**Fig 7B**). In addition, the presence of *PIK3CA* mutations drives tumor progression to mCRPC: it enhances selection for mutations associated with progression to mCRPC such as *AR*, whose critical role in resistance to therapy is well understood. The presence of *PIK3CA* mutations also enhances selection for mutations in *CTNNB1, SPOP*, and *FOXA1*. Mutations of *PIK3CA* not only function as drivers of disease but also as key intermediaries of tumor progression, contributing in ongoing ways to the oncogenic landscape of prostate cancer.

Substitutions in *CTNNB1, CUL3, KMT2C*, and *APC* enhanced selection for new mutations in *TP53* (**Fig 7C**). After oncogenic substitutions in *TP53*, selection on mutations in *PIK3CA, AKT1, PIK3CB*, and *ATM* is enhanced. An asymmetry is notable in the selective epistasis between mutations of *SPOP* and *TP53*: the presence of a mutation in *SPOP* does not highly affect selection for a subsequent *TP53* mutation. However, the presence of a mutation in *TP53* reduced the selection for a *SPOP* mutation. Therefore, we infer that during prostate cancer tumorigenesis, mutations in *TP53* typically follow mutations in *SPOP* rather than the other way around, consistent with previous evidence [78–80].

The presence of oncogenic variants in *FOXA1, PIK3CA*, and *PTEN* moderately increased selection for variants in *AR* (**Fig 7D**); the strength of selection for a mutation in *AR* is massively increased by the presence of an oncogenic variant in *KMT2D*—an epistatic change in selection for *AR* that is almost eight times the magnitude of the next-greatest positive interaction. In contrast, a mutation in *AR* increased the selection for mutations in *ATM, SPOP*, and *CTNNB1* genes. Selective epistasis corresponds to a time-ordered trajectory of somatic evolution during prostate cancer tumorigenesis.

Mutations in *PIK3CA, TP53*, and *AR* all markedly decreased the selection for mutations in *RHOA* and *CUL3*. Selection for *CUL3* mutations also decreased in the context of mutations in *SPOP*. In contrast, mutations in *SPOP* markedly increased the selection for mutation in *RHOA*, and mutations in *RHOA*, in turn, markedly increased selection for mutation in *SPOP*. No prostate cancer driver gene other than *RHOA* exhibited such reciprocated, synergistic epistasis with *SPOP* mutation. Thus, mutations in *SPOP, CUL3*, and *RHOA* are selected and spread within tumors earlier in tumorigenesis than *PIK3CA, TP53*, and (of course) *AR*.

## Discussion

Here we have shown that the trinucleotide mutation profiles—and thus COSMIC mutational signatures—underlying primary prostate tumors and mCRPC are highly similar. Dividing primary tumors into “low-grade” and “high-grade” tumors according to the Grade Groups, gene-level mutation rates increased as the tumors progressed from primary to mCRPC, with high-grade primary tumors having a higher overall mutation rate compared to low-grade primary tumors [81]. Analyzing step-specific scaled selection coefficients on recurrent mutations, we identified genes and mutations that are selected in the initiation and progression of prostate cancer from organogenesis to primary tumors to mCRPC, respectively. Annotation and depiction of the SPOP protein structure with the strength of selection on single-nucleotide driver mutations demonstrate that the strongest selection is for disruptive mutations within the binding groove; the variants with the largest scaled selection coefficient were those that would most effectively occlude the BRD-3 ligand (*i*.*e*. from a bulky R-group to a small R-group). Furthermore, analyses of epistasis support previously reported gene-gene interactions, especially interactions between *SPOP* and its epistatic pairs such as *PIK3CA*.

Within the tumor evolution from prostate organogenesis to low-grade tumors, mutations in two genes that have important roles in prostate cancer, *SPOP* and *CUL3*, were strongly selected. *SPOP* is a well-known driver that is found to be frequently mutated in prostate cancer tumors [22]. *SPOP* has two domains—a MATH domain that binds to substrates and mediates ubiquitination and subsequent degradation, and a BTB domain that assembles with CUL3, forming part of a E3 ubiquitin ligase [82]. Mutations in *CUL3* and *SPOP* have been shown to be mutually exclusive [26]. Based on our analysis, this mutual exclusivity can be quantified as negative epistatic interaction of selection on variants of these two genes.

Our analysis of selective epistasis revealed that *PTEN* mutations increase selection for mutations in both *PI3KCA* and *AR*. Two major drivers of prostate tumorigenesis, overactivation of PI3K/mTOR signaling and dysregulation of the AR signaling pathway, have been shown to negatively regulate one another through reciprocal feedback [83]. Some studies have implicated PTEN loss as a potential cause of castration resistance [84,85]. Transcriptomic analysis of PTEN-deficient tumors reveals that PI3K pathway activation suppresses AR signaling, a mechanism that may contribute to intrinsic resistance to androgen deprivation therapy. This reciprocal feedback loop enables tumor cell survival when either pathway is inhibited alone, explaining the limited efficacy of monotherapies. In contrast, combined inhibition of both PI3K and AR pathways results in tumor regression in preclinical models. Our epistatic analysis supports these findings that co-activation of PI3K and AR signaling confers a selective advantage in PTEN-null prostate cancers, underscoring the therapeutic rationale for combined pathway targeting in this molecular subtype.

*SPOP* mutations have long been thought to be early events in prostate cancer tumorigenesis [22,86]. By dividing the primary prostate tumors into “low-grade” and “high-grade” tumors—based on their Grade Groups—we showed that different genes and variants are selected for and responsible for advancing tumorigenesis at different stages of tumor evolution. Most *SPOP* mutations seem to be highly selected for in mainly the primary tumors that were a low-Grade Group. *SPOP* mutant-positive prostate tumors were analyzed and *SPOP* mutations were found to be early events that preceded any other *SPOP*-associated mutations or deletions [86]. Previous studies have shown that *TP53* mutations tend to present later and be associated with more advanced localized prostate tumors [15,31,87,88]. *SPOP* mutations and *TP53* alterations have been found to be strongly mutually exclusive [89], indicating that they represent different molecular subtypes of PC. In our analysis, the presence of mutations in *SPOP* had little effect on the selection of *TP53* mutations, while mutations in *TP53* resulted in negative epistatic change in selection for *SPOP* mutations. Furthermore, our epistasis analysis demonstrated that a mutation in *KMT2C* increases the selection for a mutation in *TP53*, with little change in selection when the order of mutations is reversed. This epistatic interaction between *TP53* and *KMT2C* may explain the previously demonstrated co-occurring relationship between *TP53* and *KMT2C* in prostate tumors [89].

The several other epistatic interactions involving *AR* are a testament to its prominent role in mCRPC. Selection for mutations in *AR* after a mutation in histone-lysine N-methyltransferase 2D (*KMT2D*, sometimes known as *MLL2*) is extremely high. Epigenetic involvement in prostate cancer tumorigenesis and metastasis is a relatively new aspect of prostate cancer research, but *KMT2D* has an established positive correlation with *AR. KMT2D* expression is correlated with not only *AR*, but also with expression of the AR-downstream target CAMKK2 [90]. KMT2D and WDR5 form a complex that increases *AR* mRNA transcription in the presence of the tyrosine kinase ACK1, which phosphorylates histone H4 upstream of the AR transcription start site [91]. Elucidating epigenetic control of *AR* may play an important role in future therapeutic options for mCRPC.

Building on the observed patterns of synergistic epistasis, epistatic interactions have important implications for disease progression. Notably, seven of the 36 synergistic pairs involved mutations associated with progression from low-grade or high-grade to mCRPC, suggesting that positive epistasis among specific driver mutations can play a critical role in enabling late-stage progression to therapeutic resistance. These epistatic interactions not only reflect temporally ordered mutational dependencies in human prostate cancer, but also reveal how the evolutionary landscape of prostate cancer is structured by interdependent selective advantages (**Fig 8**), identifying the temporal sequence and interdependent effects of key mutations. Inference of selective epistasis provides insight into both the temporal sequence and functional dependencies of somatic mutations during prostate cancer evolution. It has broad implications for understanding cancer progression and for identifying stage-specific therapeutic vulnerabilities and mutational biomarkers for precision therapeutics.

**Fig 8.**
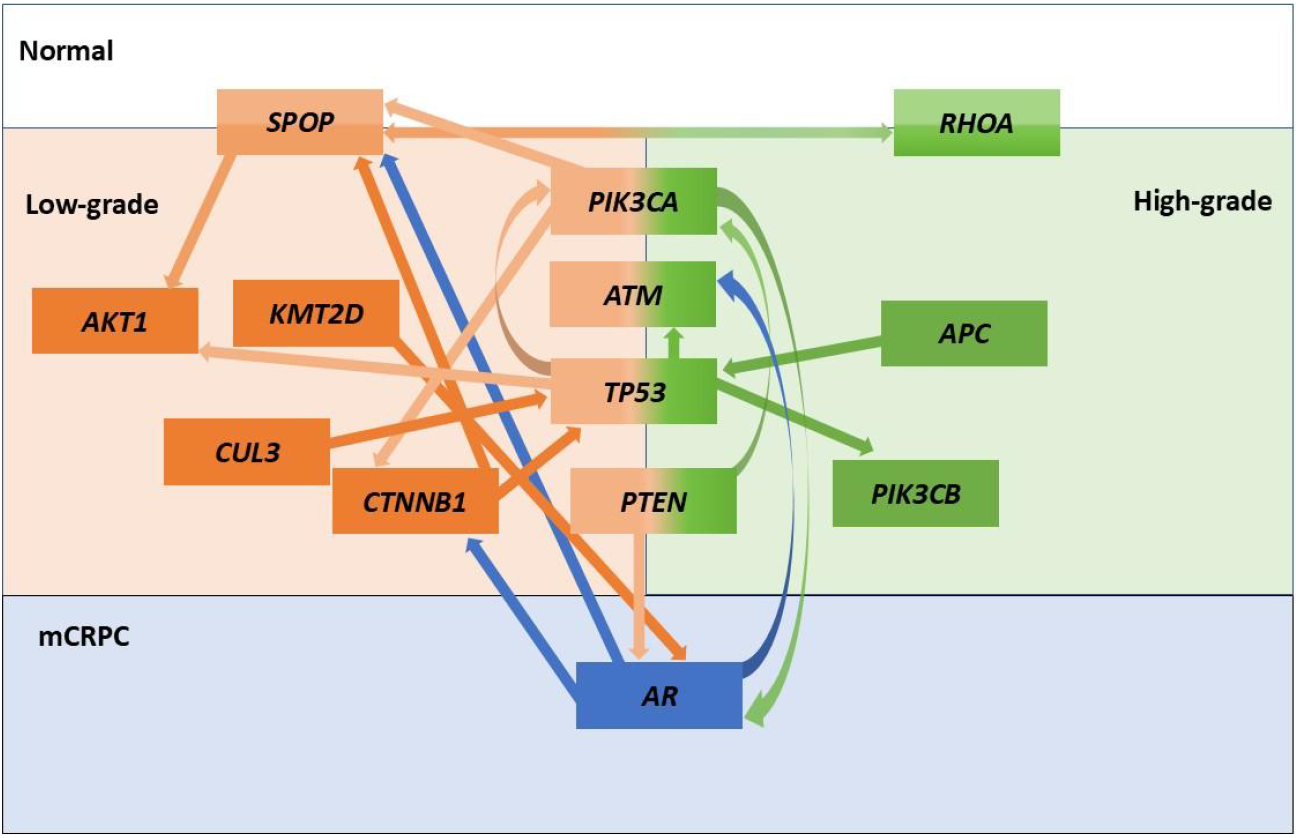
Data-driven conceptual model of some of the complex interactions among driver mutations that shape the evolution of prostate cancer. Mutations in *SPOP* are strongly selected during the transition from prostate organogenesis to low-grade tumors. In low-grade disease, *SPOP* mutations increase selection for AKT1, and in high-grade disease, they exhibit synergistic selection with mutations of *RHOA*. Mutations in *CUL3* and *CTNNB1* enhance selection for *TP53* mutations, which in turn increases selection for mutations of *PIK3CA* and *ATM*. Mutant *TP53* also enhances selection for mutations of *PIK3CB*, a gene associated—along with *RHOA* and *APC*—with high-grade tumor evolution. In the metastatic castration-resistant prostate cancer context, *PIK3CA* and *PTEN* mutations enhance selection for *AR* resistance mutations in the context of metastatic castrate-resistant prostate cancer (mCRCP), which in turn increase selection for *SPOP, CTNNB1*, and *ATM*—genes typically selected at earlier stages. *KMT2D* mutations are primarily selected for in the context of low-grade prostate cancer, but in the context of mCRPC, they increase selection for *AR* resistance mutations.

A major limitation of our analysis is that we have only been able to analyze single-nucleotide variants. There are additional genomic processes involved with prostate cancer tumorigenesis, including (1) multiple-nucleotide variants; (2) CNAs including amplifications and deletions; (3) gene fusions such as *TMPRSS2-ERG*; and (4) epigenetic changes including histone methylation and acetylation. Future research may expand our capability to analyze CNAs, which should enhance our understanding of important genes in PCs and other cancers. Also, grade heterogeneity, admixture of low- and high-grade patterns within individual tumor samples, may reduce our power to detect differences. However, the substantial differences observed in this study are unlikely to be compromised by this variability, as they remain significant despite the substantial Grade Group heterogeneity that is especially characteristic of Grade Groups 2 and 3.

A distinction among primary prostate tumors other than “low-grade” and “high-grade” could yield informative insights into prostate cancer progression and tumor evolution. However, Grade Group 1/2 prostate cancer and Grade Group 3/4/5 prostate cancer show the most striking differentiation of prognosis based on primary tumor histopathology. Our classification of genes as under selection during development into low-grade primary tumors, high-grade primary tumors, and mCRPC is only as useful as these categorizations of tumors coincide with their evolutionary and developmental biology. This differentiation of prostate cancer tumors provides a framework for further interpretation of subsequent analyses of epistatic interactions. Importantly, this differentiation does yield differences in selection coefficients inferred across genes, indicating some relation to somatic genetic evolution as well as prognosis.

Cancer effect sizes provide a metric for a gene or a somatic variant’s contribution towards tumorigenesis, but do not explain the underlying biological mechanism of selection. To fully interpret these results, cancer effect sizes should be further investigated with complementary analyses. For example, our structural analyses with *SPOP* and *AR* revealed that selected mutations occurred within the ligand binding domains of the genes, and that the most strongly selected *SPOP* variants correspond to substantial stereochemical alterations at key residues. Such remarkable findings warrant further functional investigations via traditional methods in biochemistry and molecular biology that can further establish their mechanistic roles in pathogenesis. Indeed, *SPOP* mutations differ in their biological consequences across cancers—impairing BET degradation in prostate cancer, but enhancing degradation in endometrial cancer— leading to contrasting responses to BET inhibitors [92]. Finally, broader factors such as diet and other environmental exposures can influence tumor growth [93,94] and may also have an impact on cancer effects and epistatic interactions.

In the future, cancer effect sizes can be used in conjunction with translational and clinical research to guide treatment via precision medicine, not only to treat existing mutations within the tumor, but also predict which mutations are likely to arise because of this treatment. For example, using CES analysis, *KRAS* G12C targeted therapy in lung tumors was determined to most likely give rise to *BRAF* V600E mutations [95]. In addition, knowing the evolutionary trajectory of mutations within a disease process, such as the pathogenesis of PC, will provide suggestions as to which genes to target first.

Our analysis revealed the evolutionary genetic differences in mutation and selection arising between low-grade, high-grade, and mCRPC. Gene mutation rates and selection change during progression from low-grade to high-grade and mCRPC. Strong selective epistasis among driver genes offers a compelling explanation for these dynamics. We show that *SPOP* is involved in early tumorigenesis of PC—not only by demonstration of the high selection coefficients of *SPOP* mutations in low-grade prostate tumors, but also by epistatic analysis showing that mutations in *SPOP* increase the selection for other prostate cancer driver mutations. The epistatic interactions between the many genes involved in prostate cancer tumorigenesis support a comprehensive hypothesis for its evolutionary trajectory. This quantitative analysis of the evolutionary trajectory of mutations in prostate cancer reveals the key somatic mutations underlying oncogenesis and mCRPC progression and has the potential to inform and guide future precision treatments.

## Supporting information

**S1 Table.** Mutation rates of five genes that play distinct roles in development, progression, or treatment resistance

| Gene | Gene mutation rate |  |  |  |
| --- | --- | --- | --- | --- |
|  | Organogenesis →<br>primary tumor | Organogenesis →<br>low-grade tumor | Organogenesis →<br>high-grade tumor | Organogenesis →<br>mCRPC |
| *FOXA1* | $7.5 \times 10^{-7}$ | $6.5 \times 10^{-7}$ | $8.3 \times 10^{-7}$ | $2.2 \times 10^{-6}$ |
| *AR* | $6.8 \times 10^{-7}$ | $7.2 \times 10^{-7}$ | $6.6 \times 10^{-7}$ | $1.5 \times 10^{-6}$ |
| *TP53* | $3.4 \times 10^{-7}$ | $2.5 \times 10^{-7}$ | $4.2 \times 10^{-7}$ | $1.2 \times 10^{-6}$ |
| *SPOP* | $3.5 \times 10^{-7}$ | $2.9 \times 10^{-7}$ | $4.1 \times 10^{-7}$ | $1.0 \times 10^{-6}$ |
| *PIK3CA* | $3.3 \times 10^{-7}$ | $3.0 \times 10^{-7}$ | $3.6 \times 10^{-7}$ | $8.0 \times 10^{-7}$ |

**S2 Table.** Positive epistatic interactions among driver genes across prostate cancer progression.

| Gene /<br>Gene pair<sup>a</sup> | Scaled selection coefficient /<br>Epistatic scaled selection coefficient | Effect on progression |
| --- | --- | --- |
| *CTNNB1* | 5,598 |  |
| *KMT2D*→*CTNNB1* | 166,700 | Low-grade → Low-grade |
| *AR* | 2,794 |  |
| *KMT2D*→*AR* | 79,546 | Low-grade → mCRPC |
| *AKT1* | 4,183 |  |
| *FOXA1*→*AKT1* | 73,767 | Low-grade → Low-grade |
| *RHOA* | 4,641 |  |
| *SPOP*→*RHOA* | 70,195 | Low-grade → High-grade |
| *AKT1* | 4,193 |  |
| *CTNNB1*→*AKT1* | 59,032 | Low-grade → Low-grade |
| *CTNNB1* | 5,312 |  |
| *PIK3CA*→*CTNNB1* | 49,226 | Low-grade → Low-grade |
| *ATM* | 1,688 |  |
| *FOXA1*→*ATM* | 45,276 | Low-grade → Low-grade |
| *ATM* | 1,122 |  |
| *AR*→*ATM* | 42,830 |  |
| *PTEN* | 759 |  |
| *PIK3CB*→*PTEN* | 37,215 |  |
| *CTNNB1* | 5,326 |  |
| *FOXA1*→*CTNNB1* | 35,308 | Low-grade → Low-grade |
| *SPOP* | 8,703 |  |
| *RHOA*→*SPOP* | 25,874 |  |
| *SPOP* | 8,503 |  |
| *PIK3CA*→*SPOP* | 25,213 | Low-grade → Low-grade |
| *SPOP* | 8,192 |  |
| *AR*→*SPOP* | 21,231 |  |
| *PTEN* | 800 |  |
| *CUL3*→*PTEN* | 21,162 | Low-grade → High-grade |
| *PIK3CA* | 2,963 |  |
| *PTEN*→*PIK3CA* | 16,191 |  |
| *AR* | 2,804 |  |
| *PTEN*→*AR* | 9,987 | High-grade → mCRPC |
| *SPOP* | 8,606 |  |
| *CTNNB1*→*SPOP* | 7,811 | Low-grade → Low-grade |
| *FOXA1* | 796 |  |
| *PIK3CA*→*FOXA1* | 7,583 | Low-grade → Low-grade |
| *CTNNB1* | 5,519 |  |
| *AR*→*CTNNB1* | 7,155 |  |
| *ATM* | 1,749 |  |
| *TP53*→*ATM* | 7,117 | Low-grade → Low-grade |
| *TP53* | 1,101 |  |
| *APC*→*TP53* | 6,643 |  |
| *PIK3CB* | 3,805 |  |
| *PTEN*→*PIK3CB* | 6,386 | High-grade → High-grade |
| *AR* | 2,790 |  |
| *PIK3CA*→*AR* | 6,366 | Low-grade → mCRPC |
| *TP53* | 1,093 |  |
| *KMT2C*→*TP53* | 6,080 | Low-grade → Low-grade |
| *AKT1* | 4,750 |  |
| *SPOP*→*AKT1* | 5,234 | Low-grade → Low-grade |
| *AR* | 2,795 |  |
| *FOXA1*→*AR* | 4,706 | Low-grade → mCRPC |
| *TP53* | 1,103 |  |
| *CUL3*→*TP53* | 2,739 | Low-grade → Low-grade |
| *PIK3CB* | 3,680 |  |
| *TP53*→*PIK3CB* | 2,481 | Low-grade → High-grade |
| *AKT1* | 4,844 |  |
| *TP53*→*AKT1* | 2,055 | Low-grade → Low-grade |
| *TP53* | 1,091 |  |
| *CTNNB1*→*TP53* | 1,375 | Low-grade → Low-grade |
| *FOXA1* | 821 |  |
| *SPOP*→*FOXA1* | 959 | Low-grade → Low-grade |
| *PIK3CA* | 2,975 |  |
| *TP53*→*PIK3CA* | 918 | Low-grade → Low-grade |
| *FOXA1* | 860 |  |
| *ATM*→*FOXA1* | 463 | Low-grade → Low-grade |
| *PIK3CA* | 3,034 |  |
| *FOXA1*→*PIK3CA* | 254 | Low-grade → Low-grade |
| *PTEN* | 836 |  |
| *SPOP*→*PTEN* | 240 | Low-grade → High-grade |
| *TP53* | 1,109 |  |
| *FOXA1*→*TP53* | 64 | Low-grade → Low-grade |

<sup>a</sup> A→B corresponds to selection for mutations in gene B given a mutation in gene A.

**S1 Fig.**
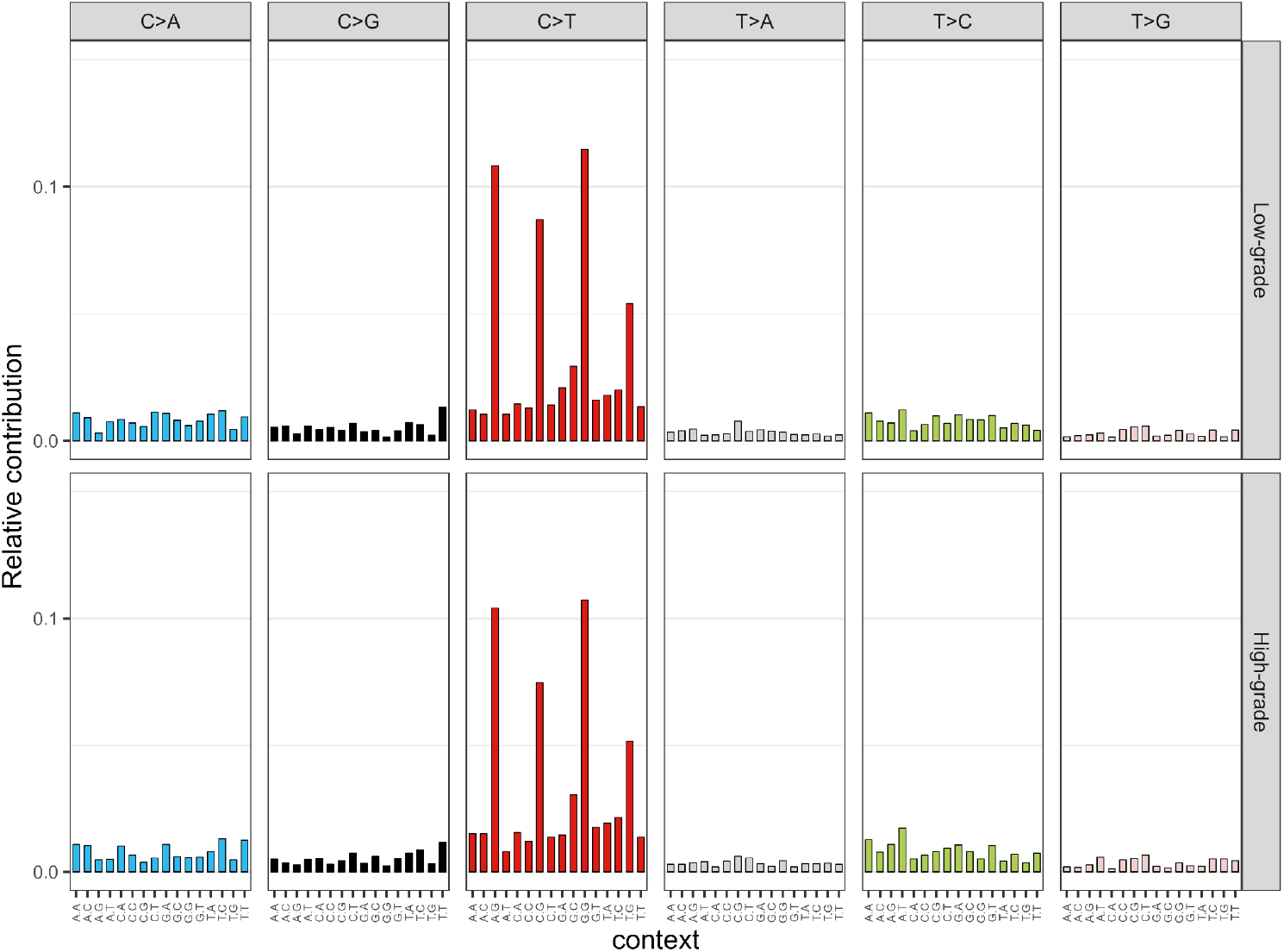
Percentage of single-nucleotide somatic variants within each trinucleotide context in low-grade and high-grade primary tumors, in an age-matched patient cohort (*n* = 318).

**S2 Fig.**
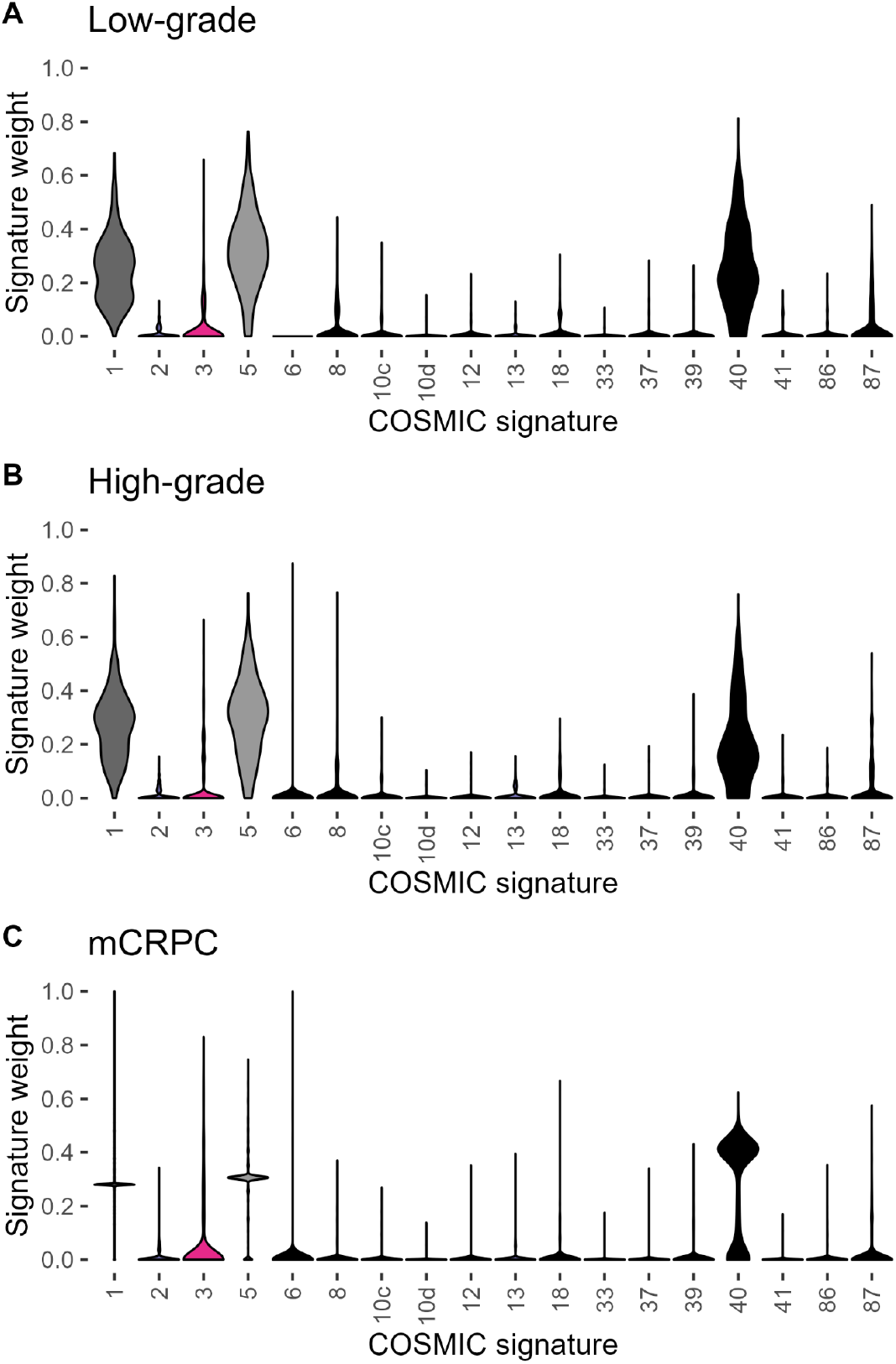
Distributions of the weights assigned to COSMIC signatures 1 (age-associated, dark gray), 2, 3 (coral, defective homologous recombination-based DNA damage repair), 5 (age-associated, light gray), 6, 8, 10c, 10d, 12, 13, 18, 33, 37, 39, 40 (black, unknown), 41, 86, and 87 in (**A**) low-grade and (**B**) high-grade primary tumors and (**C**) metastatic castrate-resistant prostate cancer tissue. Signatures not shown because they contributed no weight were 4, 7a, 7b, 7c, 7d, 9, 10a, 10b, 11, 14, 15, 16, 17a, 17b, 19, 20, 21, 22, 23, 24, 25, 26, 28, 29, 3o, 31, 32, 34, 35, 36, 38, 42, 44, 84, 85, 88, 89, 90, 91, 92, 93, and 94.

**S3 Fig.**
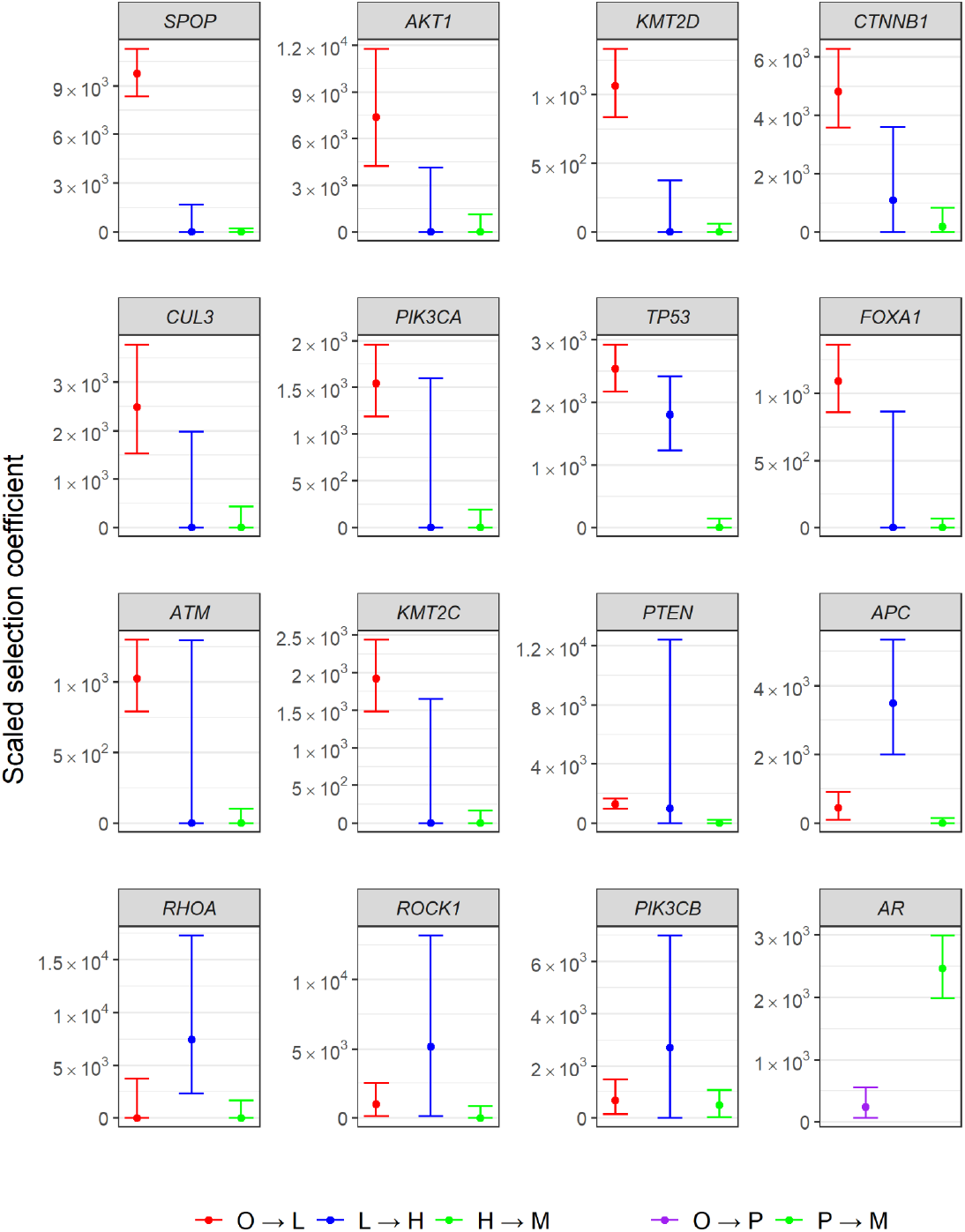
Gene-level estimates and 95% confidence intervals for scaled selection coefficients on somatic variants based on the sequential ordering in the stage-specific selection calculation, as a single trajectory from prostate organogenesis to low-grade (O→L, red) to high-grade (L→H, blue) prostate cancer to metastatic castrate-resistant prostate cancer (H→M, green) in oncogenic sites of 16 genes known to act as drivers in prostate cancer tumorigenesis and mCRPC. Estimates of underlying mutation rates for *AR* did not accommodate the disfavored linear model, and could only be estimated by combining low- and high-grade primary tumors compared to metastatic tissues in a two-stage analysis (Organogenesis → primary tumor (purple) → mCRPC (green).

## References

1. Bray F, Laversanne M, Sung H, Ferlay J, Siegel RL, Soerjomataram I, et al. Global cancer statistics 2022: GLOBOCAN estimates of incidence and mortality worldwide for 36 cancers in 185 countries. CA Cancer J Clin. 2024;74: 229–263.

2. James ND, Tannock I, N’Dow J, Feng F, Gillessen S, Ali SA, et al. The Lancet Commission on prostate cancer: planning for the surge in cases. Lancet. 2024;403: 1683–1722.

3. Frendl DM, FitzGerald G, Epstein MM, Allison JJ, Sokoloff MH, Ware JE. Predicting the 10-year risk of death from other causes in men with localized prostate cancer using patient-reported factors: Development of a tool. PLoS One. 2020;15: e0240039.

4. Hamdy FC, Donovan JL, Lane JA, Metcalfe C, Davis M, Turner EL, et al. Fifteen-Year Outcomes after Monitoring, Surgery, or Radiotherapy for Prostate Cancer. N Engl J Med. 2023;388: 1547–1558.

5. Siegel RL, Miller KD, Jemal A. Cancer statistics, 2018. CA: A Cancer Journal for Clinicians. 2018;68: 7–30.

6. Vogelstein B, Kinzler KW. The multistep nature of cancer. Trends Genet. 1993;9: 138–141.

7. Sowalsky AG, Kissick HT, Gerrin SJ, Schaefer RJ, Xia Z, Russo JW, et al. Gleason Score 7 Prostate Cancers Emerge through Branched Evolution of Clonal Gleason Pattern 3 and 4. Clin Cancer Res. 2017;23: 3823–3833.

8. Boutros PC, Fraser M, Harding NJ, de Borja R, Trudel D, Lalonde E, et al. Spatial genomic heterogeneity within localized, multifocal prostate cancer. Nat Genet. 2015;47: 736–745.

9. Cooper CS, Eeles R, Wedge DC, Van Loo P, Gundem G, Alexandrov LB, et al. Analysis of the genetic phylogeny of multifocal prostate cancer identifies multiple independent clonal expansions in neoplastic and morphologically normal prostate tissue. Nat Genet. 2015;47: 367–372.

10. Buhigas C, Warren AY, Leung W-K, Whitaker HC, Luxton HJ, Hawkins S, et al. The architecture of clonal expansions in morphologically normal tissue from cancerous and non-cancerous prostates. Mol Cancer. 2022;21: 183.

11. VanderWeele DJ, Brown CD, Taxy JB, Gillard M, Hatcher DM, Tom WR, et al. Low-grade prostate cancer diverges early from high grade and metastatic disease. Cancer Sci. 2014;105: 1079–1085.

12. Taylor RA, Fraser M, Livingstone J, Espiritu SMG, Thorne H, Huang V, et al. Germline BRCA2 mutations drive prostate cancers with distinct evolutionary trajectories. Nat Commun. 2017;8: 1–10.

13. Berman DM, Lee AY, Lesurf R, Patel PG, Ebrahimizadeh W, Bayani J, et al. Multimodal Biomarkers That Predict the Presence of Gleason Pattern 4: Potential Impact for Active Surveillance. J Urol. 2023;210: 257–271.

14. Brastianos HC, Murgic J, Salcedo A, Chua MLK, Meng A, Fraser M, et al. Determining the Impact of Spatial Heterogeneity on Genomic Prognostic Biomarkers for Localized Prostate Cancer. Eur Urol Oncol. 2022;5: 362–365.

15. Espiritu SMG, Liu LY, Rubanova Y, Bhandari V, Holgersen EM, Szyca LM, et al. The Evolutionary Landscape of Localized Prostate Cancers Drives Clinical Aggression. Cell. 2018;173: 1003–1013.e15.

16. Quigley DA, Dang HX, Zhao SG, Lloyd P, Aggarwal R, Alumkal JJ, et al. Genomic Hallmarks and Structural Variation in Metastatic Prostate Cancer. Cell. 2018;174: 758–769.e9.

17. Robinson D, Van Allen EM, Wu Y-M, Schultz N, Lonigro RJ, Mosquera J-M, et al. Integrative clinical genomics of advanced prostate cancer. Cell. 2015;161: 1215–1228.

18. Gundem G, Van Loo P, Kremeyer B, Alexandrov LB, Tubio JMC, Papaemmanuil E, et al. The evolutionary history of lethal metastatic prostate cancer. Nature. 2015;520: 353–357.

19. Beltran H, Prandi D, Mosquera JM, Benelli M, Puca L, Cyrta J, et al. Divergent clonal evolution of castration-resistant neuroendocrine prostate cancer. Nat Med. 2016;22: 298–305.

20. Fraser M, Sabelnykova VY, Yamaguchi TN, Heisler LE, Livingstone J, Huang V, et al. Genomic hallmarks of localized, non-indolent prostate cancer. Nature. 2017;541: 359–364.

21. Kumar A, Coleman I, Morrissey C, Zhang X, True LD, Gulati R, et al. Substantial interindividual and limited intraindividual genomic diversity among tumors from men with metastatic prostate cancer. Nat Med. 2016;22: 369–378.

22. Barbieri CE, Baca SC, Lawrence MS, Demichelis F, Blattner M, Theurillat J-P, et al. Exome sequencing identifies recurrent SPOP, FOXA1 and MED12 mutations in prostate cancer. Nat Genet. 2012;44: 685–689.

23. Spratt DE, Zumsteg ZS, Feng FY, Tomlins SA. Translational and clinical implications of the genetic landscape of prostate cancer. Nat Rev Clin Oncol. 2016;13: 597–610.

24. Wedge DC, Gundem G, Mitchell T, Woodcock DJ, Martincorena I, Ghori M, et al. Sequencing of prostate cancers identifies new cancer genes, routes of progression and drug targets. Nat Genet. 2018;50: 682–692.

25. Cancer Genome Atlas Research Network. The Molecular Taxonomy of Primary Prostate Cancer. Cell. 2015;163: 1011–1025.

26. Armenia J, Wankowicz SAM, Liu D, Gao J, Kundra R, Reznik E, et al. The long tail of oncogenic drivers in prostate cancer. Nat Genet. 2018;50: 645–651.

27. Gerhauser C, Favero F, Risch T, Simon R, Feuerbach L, Assenov Y, et al. Molecular Evolution of Early-Onset Prostate Cancer Identifies Molecular Risk Markers and Clinical Trajectories. Cancer Cell. 2018;34: 996–1011.e8.

28. van Dessel LF, van Riet J, Smits M, Zhu Y, Hamberg P, van der Heijden MS, et al. The genomic landscape of metastatic castration-resistant prostate cancers reveals multiple distinct genotypes with potential clinical impact. Nat Commun. 2019;10: 5251.

29. Rebello RJ, Oing C, Knudsen KE, Loeb S, Johnson DC, Reiter RE, et al. Prostate cancer. Nat Rev Dis Primers. 2021;7: 9.

30. Grasso CS, Wu Y-M, Robinson DR, Cao X, Dhanasekaran SM, Khan AP, et al. The mutational landscape of lethal castration-resistant prostate cancer. Nature. 2012;487: 239–243.

31. Fraser M, Livingstone J, Wrana JL, Finelli A, He HH, van der Kwast T, et al. Somatic driver mutation prevalence in 1844 prostate cancers identifies ZNRF3 loss as a predictor of metastatic relapse. Nat Commun. 2021;12: 6248.

32. Shorning BY, Dass MS, Smalley MJ, Pearson HB. The PI3K-AKT-mTOR Pathway and Prostate Cancer: At the Crossroads of AR, MAPK, and WNT Signaling. Int J Mol Sci. 2020;21. doi:10.3390/ijms21124507

33. Pearson HB, Li J, Meniel VS, Fennell CM, Waring P, Montgomery KG, et al. Identification of Pik3ca Mutation as a Genetic Driver of Prostate Cancer That Cooperates with Pten Loss to Accelerate Progression and Castration-Resistant Growth. Cancer Discovery. 2018. pp. 764–779. doi:10.1158/2159-8290.cd-17-0867

34. Engelman JA. Targeting PI3K signalling in cancer: opportunities, challenges and limitations. Nat Rev Cancer. 2009;9: 550–562.

35. Hodgson MC, Astapova I, Cheng S, Lee LJ, Verhoeven MC, Choi E, et al. The androgen receptor recruits nuclear receptor CoRepressor (N-CoR) in the presence of mifepristone via its N and C termini revealing a novel molecular mechanism for androgen receptor antagonists. J Biol Chem. 2005;280: 6511–6519.

36. Hsieh C-L, Botta G, Gao S, Li T, Van Allen EM, Treacy DJ, et al. PLZF, a tumor suppressor genetically lost in metastatic castration-resistant prostate cancer, is a mediator of resistance to androgen deprivation therapy. Cancer Res. 2015;75: 1944–1948.

37. van der Steen T, Tindall DJ, Huang H. Posttranslational modification of the androgen receptor in prostate cancer. Int J Mol Sci. 2013;14: 14833–14859.

38. Antonarakis ES, Armstrong AJ, Dehm SM, Luo J. Androgen receptor variant-driven prostate cancer: clinical implications and therapeutic targeting. Prostate Cancer Prostatic Dis. 2016;19: 231–241.

39. Valle LF, Li J, Desai H, Hausler R, Haroldsen C, Chatwal M, et al. Somatic Tumor Next-Generation Sequencing in US Veterans With Metastatic Prostate Cancer. JAMA Netw Open. 2025;8: e259119.

40. Gerhauser C, Favero F, Risch T, Simon R, Feuerbach L, Assenov Y, et al. Molecular Evolution of Early-Onset Prostate Cancer Identifies Molecular Risk Markers and Clinical Trajectories. Cancer Cell. 2018;34: 996–1011.e8.

41. Ramazzotti D, Caravagna G, Olde Loohuis L, Graudenzi A, Korsunsky I, Mauri G, et al. CAPRI: efficient inference of cancer progression models from cross-sectional data. Bioinformatics. 2015;31: 3016–3026.

42. Cannataro VL, Gaffney SG, Townsend JP. Effect Sizes of Somatic Mutations in Cancer. J Natl Cancer Inst. 2018;110: 1171–1177.

43. Dasari K, Somarelli JA, Kumar S, Townsend JP. The somatic molecular evolution of cancer: Mutation, selection, and epistasis. Prog Biophys Mol Biol. 2021;165: 56–65.

44. Wilkins JF, Cannataro VL, Shuch B, Townsend JP. Analysis of mutation, selection, and epistasis: an informed approach to cancer clinical trials. Oncotarget. 2018;9: 22243–22253.

45. Pairwise and higher-order epistatic effects among somatic cancer mutations across oncogenesis. Mathematical Biosciences. 2023;366: 109091.

46. Abida W, Cyrta J, Heller G, Prandi D, Armenia J, Coleman I, et al. Genomic correlates of clinical outcome in advanced prostate cancer. Proc Natl Acad Sci U S A. 2019;116: 11428–11436.

47. Stopsack KH, Nandakumar S, Wibmer AG, Haywood S, Weg ES, Barnett ES, et al. Oncogenic Genomic Alterations, Clinical Phenotypes, and Outcomes in Metastatic Castration-Sensitive Prostate Cancer. Clin Cancer Res. 2020;26: 3230–3238.

48. Epstein JI, Egevad L, Amin MB, Delahunt B, Srigley JR, Humphrey PA, et al. The 2014 International Society of Urological Pathology (ISUP) Consensus Conference on Gleason Grading of Prostatic Carcinoma: Definition of Grading Patterns and Proposal for a New Grading System. Am J Surg Pathol. 2016;40: 244–252.

49. Pierorazio PM, Walsh PC, Partin AW, Epstein JI. Prognostic Gleason grade grouping: data based on the modified Gleason scoring system. BJU International. 2013. pp. 753–760. doi:10.1111/j.1464-410x.2012.11611.x

50. Martincorena I, Raine KM, Gerstung M, Dawson KJ, Haase K, Van Loo P, et al. Universal Patterns of Selection in Cancer and Somatic Tissues. Cell. 2018;173: 1823.

51. Mandell JD, Cannataro VL, Townsend JP. Estimation of Neutral Mutation Rates and Quantification of Somatic Variant Selection Using cancereffectsizeR. Cancer Res. 2023;83: 500–505.

52. Blokzijl F, Janssen R, van Boxtel R, Cuppen E. MutationalPatterns: comprehensive genome-wide analysis of mutational processes. Genome Med. 2018;10: 33.

53. Cannataro VL, Mandell JD, Townsend JP. Attribution of Cancer Origins to Endogenous, Exogenous, and Preventable Mutational Processes. Mol Biol Evol. 2022;39. doi:10.1093/molbev/msac084

54. Robinson DR, Wu Y-M, Lonigro RJ, Vats P, Cobain E, Everett J, et al. Integrative clinical genomics of metastatic cancer. Nature. 2017;548: 297–303.

55. Liu K, Li X, Wang J, Wang Y, Dong H, Li J. Genetic variants in RhoA and ROCK1 genes are associated with the development, progression and prognosis of prostate cancer. Oncotarget. 2017;8: 19298–19309.

56. Lv S, Ji L, Chen B, Liu S, Lei C, Liu X, et al. Histone methyltransferase KMT2D sustains prostate carcinogenesis and metastasis via epigenetically activating LIFR and KLF4. Oncogene. 2018;37: 1354–1368.

57. Glasmacher KA, Cannataro VL, Mandell JD, Jackson M, Nic Fisk J, Townsend JP. Mutation of NOTCH1 is selected within normal esophageal tissues, yet leads to selective epistasis suppressive of further evolution into cancer. bioRxiv. 2023. p. 2023.11.03.565535. doi:10.1101/2023.11.03.565535

58. Berman HM, Westbrook J, Feng Z, Gilliland G, Bhat TN, Weissig H, et al. The Protein Data Bank. Nucleic Acids Res. 2000;28: 235–242.

59. Zhou XE, Suino-Powell KM, Li J, He Y, Mackeigan JP, Melcher K, et al. Identification of SRC3/AIB1 as a preferred coactivator for hormone-activated androgen receptor. J Biol Chem. 2010;285: 9161–9171.

60. Pettersen EF, Goddard TD, Huang CC, Couch GS, Greenblatt DM, Meng EC, et al. UCSF Chimera?A visualization system for exploratory research and analysis. Journal of Computational Chemistry. 2004. pp. 1605–1612. doi:10.1002/jcc.20084

61. Alexandrov LB, Kim J, Haradhvala NJ, Huang MN, Ng AWT, Wu Y, et al. The repertoire of mutational signatures in human cancer. Nature. 2020;578: 94–101.

62. Grossmann S, Hooks Y, Wilson L, Moore L, O’Neill L, Martincorena I, et al. Development, maturation, and maintenance of human prostate inferred from somatic mutations. Cell Stem Cell. 2021. doi:10.1016/j.stem.2021.02.005

63. Leitzmann MF, Rohrmann S. Risk factors for the onset of prostatic cancer: age, location, and behavioral correlates. Clin Epidemiol. 2012;4: 1–11.

64. Priestley P, Baber J, Lolkema MP, Steeghs N, de Bruijn E, Shale C, et al. Pan-cancer whole-genome analyses of metastatic solid tumours. Nature. 2019;575: 210–216.

65. Fujita K, Nonomura N. Role of Androgen Receptor in Prostate Cancer: A Review. World J Mens Health. 2019;37: 288–295.

66. Koushyar S, Meniel VS, Phesse TJ, Pearson HB. Exploring the Wnt Pathway as a Therapeutic Target for Prostate Cancer. Biomolecules. 2022;12. doi:10.3390/biom12020309

67. Zhuang M, Calabrese MF, Liu J, Waddell MB, Nourse A, Hammel M, et al. Structures of SPOP-substrate complexes: insights into molecular architectures of BTB-Cul3 ubiquitin ligases. Mol Cell. 2009;36: 39–50.

68. Cannataro VL, Townsend JP. Wagging the long tail of drivers of prostate cancer. PLOS Genetics. 2019. p. e1007820. doi:10.1371/journal.pgen.1007820

69. Chen CD, Welsbie DS, Tran C, Baek SH, Chen R, Vessella R, et al. Molecular determinants of resistance to antiandrogen therapy. Nat Med. 2004;10: 33–39.

70. Watson PA, Arora VK, Sawyers CL. Emerging mechanisms of resistance to androgen receptor inhibitors in prostate cancer. Nat Rev Cancer. 2015;15: 701–711.

71. Crona DJ, Whang YE. Androgen receptor-dependent and -independent mechanisms involved in prostate cancer therapy resistance. Cancers (Basel). 2017;9: 67.

72. Veldscholte J, Ris-Stalpers C, Kuiper GGJM, Jenster G, Berrevoets C, Claassen E, et al. A mutation in the ligand binding domain of the androgen receptor of human INCaP cells affects steroid binding characteristics and response to anti-androgens. Biochem Biophys Res Commun. 1990;173: 534–540.

73. Tan J, Sharief Y, Hamil KG, Gregory CW, Zang DY, Sar M, et al. Dehydroepiandrosterone activates mutant androgen receptors expressed in the androgen-dependent human prostate cancer xenograft CWR22 and LNCaP cells. Mol Endocrinol. 1997;11: 450–459.

74. Hara T, Miyazaki J-I, Araki H, Yamaoka M, Kanzaki N, Kusaka M, et al. Novel mutations of androgen receptor: a possible mechanism of bicalutamide withdrawal syndrome. Cancer Res. 2003;63: 149–153.

75. Lallous N, Volik SV, Awrey S, Leblanc E, Tse R, Murillo J, et al. Functional analysis of androgen receptor mutations that confer anti-androgen resistance identified in circulating cell-free DNA from prostate cancer patients. Genome Biol. 2016;17: 10.

76. Zhao XY, Malloy PJ, Krishnan AV, Swami S, Navone NM, Peehl DM, et al. Glucocorticoids can promote androgen-independent growth of prostate cancer cells through a mutated androgen receptor. Nat Med. 2000;6: 703–706.

77. van de Wijngaart DJ, Molier M, Lusher SJ, Hersmus R, Jenster G, Trapman J, et al. Systematic Structure-Function Analysis of Androgen Receptor Leu701 Mutants Explains the Properties of the Prostate Cancer Mutant L701H 2. J Biol Chem. 2010;285: 5097–5105.

78. Zhang J, Chen M, Zhu Y, Dai X, Dang F, Ren J, et al. SPOP Promotes Nanog Destruction to Suppress Stem Cell Traits and Prostate Cancer Progression. Dev Cell. 2019;48: 329–344.e5.

79. Barbieri CE, Demichelis F, Rubin MA. The lethal clone in prostate cancer: redefining the index. Eur Urol. 2014;66: 395–397.

80. Baca SC, Prandi D, Lawrence MS, Mosquera JM, Romanel A, Drier Y, et al. Punctuated evolution of prostate cancer genomes. Cell. 2013;153: 666–677.

81. Yamaguchi TN, Houlahan KE, Zhu H, Kurganovs N, Livingstone J, Fox NS, et al. The Germline and Somatic Origins of Prostate Cancer Heterogeneity. Cancer Discov. 2025;15: 988–1017.

82. Wei X, Fried J, Li Y, Hu L, Gao M, Zhang S, et al. Functional roles of Speckle-Type Poz (SPOP) Protein in Genomic stability. J Cancer. 2018;9: 3257–3262.

83. Carver BS, Chapinski C, Wongvipat J, Hieronymus H, Chen Y, Chandarlapaty S, et al. Reciprocal feedback regulation of PI3K and androgen receptor signaling in PTEN-deficient prostate cancer. Cancer Cell. 2011;19: 575–586.

84. Zhang W, Zhu J, Efferson CL, Ware C, Tammam J, Angagaw M, et al. Inhibition of Tumor Growth Progression by Antiandrogens and mTOR Inhibitor in a Pten-Deficient Mouse Model of Prostate Cancer. Cancer Res. 2009;69: 7466–7472.

85. Gao H, Ouyang X, Banach-Petrosky WA, Shen MM, Abate-Shen C. Emergence of Androgen Independence at Early Stages of Prostate Cancer Progression in Nkx3.1; Pten Mice. Cancer Res. 2006;66: 7929–7933.

86. Boysen G, Barbieri CE, Prandi D, Blattner M, Chae S-S, Dahija A, et al. SPOP mutation leads to genomic instability in prostate cancer. Elife. 2015;4. doi:10.7554/eLife.09207

87. Kubota Y, Shuin T, Uemura H, Fujinami K, Miyamoto H, Torigoe S, et al. Tumor suppressor gene P53 mutations in human prostate cancer. The Prostate. 1995. pp. 18–24. doi:10.1002/pros.2990270105

88. Schlomm T, Iwers L, Kirstein P, Jessen B, Köllermann J, Minner S, et al. Clinical significance of p53 alterations in surgically treated prostate cancers. Mod Pathol. 2008;21: 1371–1378.

89. Liu Z, Guo H, Zhu Y, Xia Y, Cui J, Shi K, et al. TP53 alterations of hormone-naïve prostate cancer in the Chinese population. Prostate Cancer Prostatic Dis. 2021;24: 482–491.

90. Lv S-D, Wang H-Y, Yu X-P, Zhai Q-L, Wu Y-B, Wei Q, et al. Integrative molecular characterization of Chinese prostate cancer specimens. Asian J Androl. 2020;22: 162–168.

91. Mahajan K, Malla P, Lawrence HR, Chen Z, Kumar-Sinha C, Malik R, et al. ACK1/TNK2 regulates histone H4 Tyr88-phosphorylation and AR gene expression in castration-resistant prostate cancer. Cancer Cell. 2017;31: 790–803.e8.

92. Janouskova H, El Tekle G, Bellini E, Udeshi ND, Rinaldi A, Ulbricht A, et al. Opposing effects of cancer-type-specific SPOP mutants on BET protein degradation and sensitivity to BET inhibitors. Nat Med. 2017;23: 1046–1054.

93. Fontana L, Adelaiye RM, Rastelli AL, Miles KM, Ciamporcero E, Longo VD, et al. Dietary protein restriction inhibits tumor growth in human xenograft models. Oncotarget. 2013;4: 2451–2461.

94. Cordova RA, Elbanna M, Rupert C, Orsi SA, Sommers NR, Klunk AJ, et al. Caloric Restriction Enhances the Efficacy of Antiandrogen Therapy in Prostate Cancer by Inhibiting Androgen Receptor Translation. Cancer Res. 2025; OF1–OF16.

95. Cannataro VL, Gaffney SG, Stender C, Zhao Z-M, Philips M, Greenstein AE, et al. Heterogeneity and mutation in KRAS and associated oncogenes: evaluating the potential for the evolution of resistance to targeting of KRAS G12C. Oncogene. 2018;37: 2444–2455.

